# Aldoximes serve as auxin precursors and repress phenylpropanoid metabolism in tomato

**DOI:** 10.64898/2026.05.07.723529

**Authors:** Haohao Zhao, Doosan Shin, Ethan Tucker, Keun Ho Cho, Ariel Sorg, Dake Liu, Yousong Ding, Anna K. Block, Jeongim Kim

## Abstract

Aldoximes are amino acid-derived metabolites that serve as precursors of auxins and modulate phenylpropanoid production in Arabidopsis. However, the enzymes responsible for aldoxime production in Solanaceae remain unknown. Here, we report the identification of aldoxime-producing enzymes in tomato (*Solanum lycopersicum*) and examine how altered aldoxime production affects auxin production and phenylpropanoid metabolism. Through homology-based analysis, we identified five putative CYP79 homologs in tomato, among which SlCYP79DB32 and SlCYP79DB52 exhibited aldoxime-producing activity toward multiple amino acids, including phenylalanine and tryptophan. SlCYP79DB32 and SlCYP79DB52 converted phenylalanine into phenylacetaldoxime (PAOx), whereas only SlCYP79DB52 converted tryptophan into indole-3-acetaldoxime (IAOx). Stable isotope-labeled feeding experiments revealed that IAOx and PAOx can be converted to the auxins indole-3-acetic acid (IAA) and phenylacetic acid (PAA), respectively. Consistently, tomato plants engineered to overproduce IAOx and PAOx accumulated elevated levels of IAA and PAA. These plants also accumulated lower levels of phenylpropanoids. In Brassicaceae plants such as Arabidopsis and Camelina, aldoxime accumulation represses phenylpropanoid production by promoting degradation of phenylalanine ammonia-lyase (PAL). However, aldoxime accumulation did not reduce PAL activity in tomato, suggesting an alternative mechanism in this species. Transcriptome analysis revealed extensive transcriptional reprogramming in aldoxime-overaccumulating tomato plants, including upregulation of stress– and defense-related genes. Despite the observed reduction in phenylpropanoid levels, transcript levels of most phenylpropanoid biosynthetic genes were not decreased, suggesting possible post-transcriptional regulation of this repression. Together, our findings demonstrate that aldoximes can serve as intermediates in auxin biosynthesis in tomato and reveal that aldoxime-mediated repression of phenylpropanoid metabolism extends beyond Brassicaceae.

## Introduction

Aldoximes are amino acid derivatives widely distributed in the plant kingdom ^1,2^. In many higher plants, cytochrome P450 monooxygenases of the 79 family (CYP79) catalyze aldoxime formation ^3–5^. For example, in *Arabidopsis thaliana*, AtCYP79B2 and AtCYP79B3 redundantly convert tryptophan (Trp) to indole-3-acetaldoxime (IAOx), whereas AtCYP79A2 converts phenylalanine (Phe) to phenylacetaldoxime (PAOx) ^6–8^. In sorghum, SbCYP79A61 and SbCYP79A1 produce PAOx and 4-hydroxy PAOx from Phe and tyrosine (Tyr), respectively ^3,9^. In poplar, PtCYP79D6v3 and PtCYP79D7v2 convert Trp, Phe, leucine (Leu), and isoleucine (Ile) to IAOx, PAOx, 3-methylbutyraldoxime, and 2-methylbutyraldoxime, respectively ^10^.

Aldoximes can integrate plant growth and stress responses by modulating multiple metabolic pathways. They are well recognized as precursors of defense compounds, such as cyanogenic glycosides and glucosinolates, which confer resistance to herbivores and pathogens ^1,2^. Some aldoximes or their derivatives are also emitted as volatiles, playing diverse roles in plant-environment interactions, such as pollinator attraction, herbivore repellence, and recruitment of herbivore enemies ^1,2^. In addition, aldoximes can serve as precursors of the naturally occurring auxins indole-3-acetic acid (IAA) and phenylacetic acid (PAA), which regulate plant growth, development, and stress responses ^11^. Although IAA is generally considered a more potent auxin than PAA, PAA often accumulates to higher levels than IAA in various plant species ^12^. In most plants, IAA is produced mainly through the YUCCA (YUC) pathway, which involves Tryptophan Aminotransferase of Arabidopsis (TAA) family aminotransferases and YUC family flavin-containing monooxygenases ^13,14^. PAA has been proposed to be synthesized through an analogous pathway, in which TAAs convert phenylalanine to phenylpyruvate (PPA), and YUC enzymes subsequently convert PPA to PAA ^15–17^. Besides the YUC pathway, several alternative auxin biosynthesis pathways have been proposed ^18,19^. One such pathway is the aldoxime-mediated auxin pathway, which uses IAOx and PAOx as precursors for IAA and PAA, respectively ^7,20,21^. As IAOx and PAOx are precursors of indole glucosinolates and benzyl glucosinolate, which are commonly found in Brassicaceae, this pathway was initially considered Brassicaceae-specific. However, recent studies have identified IAA production in IAOx-producing maize ^22^ and the conversion of PAOx to PAA in sorghum^9^, although the conversion steps from aldoximes to auxins remain largely unknown.

Aldoxime metabolism also influences plant growth and development through its interaction with the phenylpropanoid pathway. Phenylpropanoids are specialized metabolites derived primarily from Phe and include lignin, flavonoids, and hydroxycinnamate derivatives, which contribute to structural support, pigmentation, UV protection, and defense ^23^. In Arabidopsis, accumulation of aromatic and aliphatic aldoximes represses phenylpropanoid biosynthesis ^22,24–27^. Mechanistically, this repression is mediated by increased degradation of phenylalanine ammonia-lyase (PAL), the entry-point enzyme of the phenylpropanoid pathway. Aldoxime accumulation activates the transcription of Kelch repeat F-box (KFB) proteins that target PAL for ubiquitin-dependent proteasomal turnover ^24,25,28^. However, whether this regulatory mechanism is conserved beyond the Brassicaceae remains unknown.

The Solanaceae family includes economically important crops such as potato, tomato, pepper, eggplant, and tobacco ^29^. Some aldoxime derivatives have been detected in stressed eggplant (*Solanum melongena*): 2-methylpropanal-*O*-methyl oxime and 3-methylbutanal-*O*-methyl oxime were emitted in spider mite-infested eggplant leaves, while 2-methylbutanal-*O*-methyl oxime was emitted in clean and mechanically damaged leaves ^30^. Despite the significance of aldoximes, auxins, and phenylpropanoids for plant growth and survival, the identity of aldoxime-producing enzymes is unknown in Solanaceae.

As a model species in the Solanaceae family and one of the most widely consumed crops worldwide, tomato (*Solanum lycopersicum*) provides an excellent system for investigating aldoxime metabolism ^31,32^. Here, we identified five putative CYP79-encoding genes in the tomato genome and investigated whether they can catalyze aldoxime formation and the metabolic connectivity of aldoximes with the production of auxins and phenylpropanoids. Our data revealed that tomato CYP79s generate PAOx, IAOx, and additional aliphatic aldoximes, and that PAOx and IAOx are converted to the auxins PAA and IAA, respectively. Furthermore, elevated aldoxime production in vivo strongly represses phenylpropanoid metabolism. Together, these findings establish aldoximes as intermediates in auxin biosynthesis and reveal a metabolic connection between aldoxime accumulation and phenylpropanoid repression in tomato.

## Result

### Identification of CYP79 enzymes in tomato

To identify CYP79 enzymes that can convert amino acids into aldoximes in tomato (Fig. 1A), we searched the tomato genome for cytochrome P450 (CYP)-encoding genes and identified 283 putative CYP genes (Fig. 1B; Supplementary Fig. S1). Among them, five tomato CYP79 homologs, SlCYP79DB32 (Solyc08g083450), SlCYP79DB43 (Solyc04g025410), SlCYP79DB46 (Solyc04g005370), SlCYP79DB47 (Solyc04g005360), and SlCYP79DB52 (Solyc12g005170), named following the revised CYP79 nomenclature ^33^, clustered with functionally characterized CYP79 enzymes from diverse species (Fig. 1B). A multiple sequence alignment of tomato CYP79 candidates with characterized CYP79 enzymes showed that the tomato sequences contain conserved structural features typical of plant A-type cytochrome P450s ^34^, including the N-terminal proline-rich PPGP/PPXP motif and a conserved heme-binding region containing the invariant catalytic cysteine residue. In addition, they share sequence features previously described in CYP79 enzymes, including an NP substitution at the position corresponding to the conserved oxygen-binding motif (AGxDT) of typical P450s and a PERH variant of the K-helix PERF motif ^34–36^ (Supplementary Fig. S2). According to transcriptomic data from the CoNekT database ^37^, these genes showed low transcript abundance across most organs, with relatively higher expression restricted to specific organs or developmental stages, indicating organ– and stage-dependent expression patterns (Fig. 1C).

**Figure 1.**
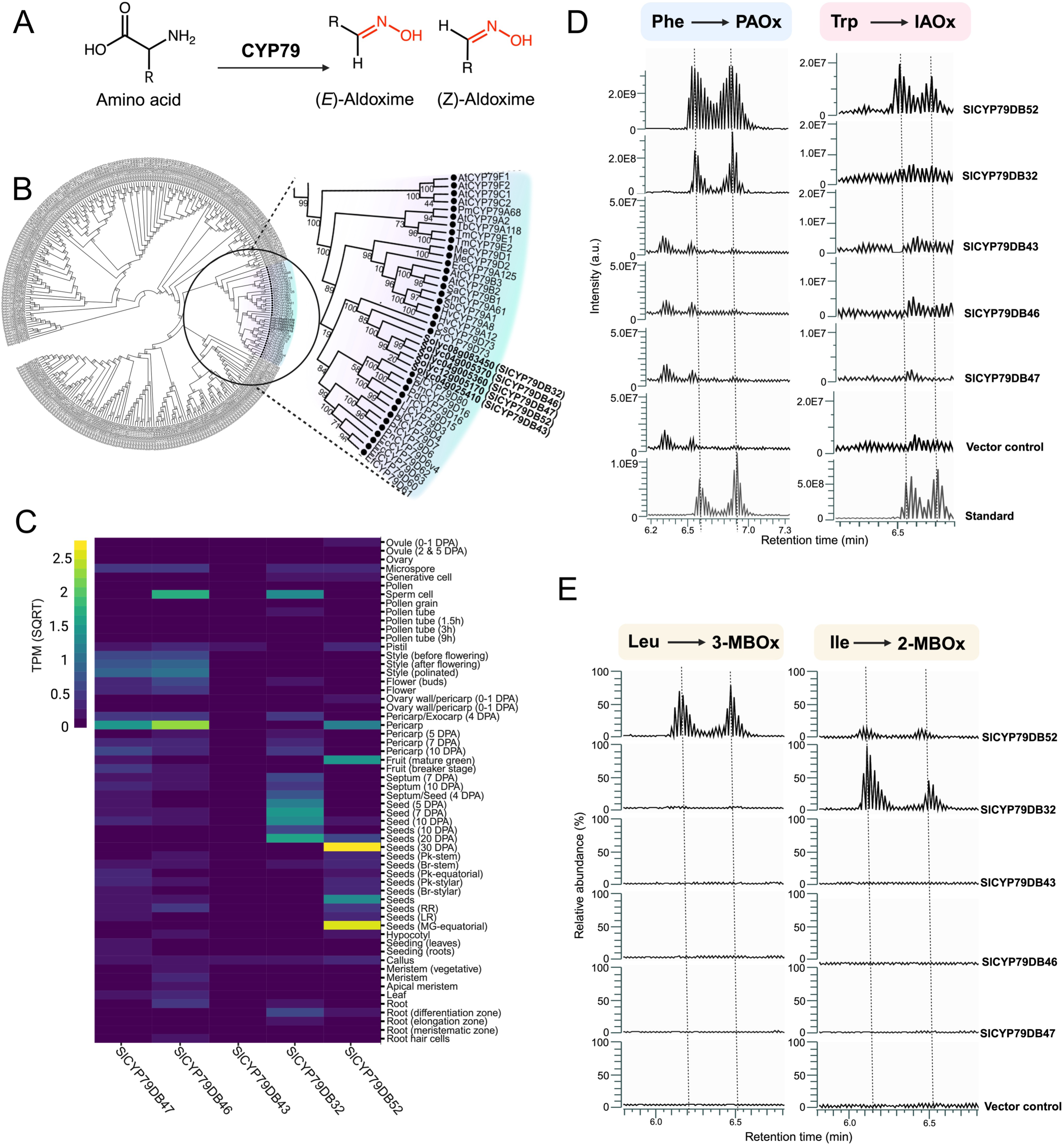
Identification and in vitro enzyme activity assay of tomato CYP79 homologs. (A) Schematic representation of the catalytic activity of CYP79 enzymes, which convert amino acids into (*E*)– and (*Z*)-aldoxime isomers. (B) Phylogenetic analysis of the plant CYP79 family, with the inset highlighting a distinct clade containing five *Solanum lycopersicum* CYP79 (SlCYP79s) clustered with characterized homologs from other species. (C) Heatmap displaying *SlCYP79* gene expression profiles across a broad range of tomato organs and developmental stages. Expression values are presented as the square root of transcripts per million, derived from the CoNekT database ^37^. (D, E) In vitro enzymatic activity assays of recombinant SlCYP79 proteins. Recombinant SlCYP79DB32, 43, 46, 47, and 52 proteins expressed in *E. coli* were incubated with amino acid substrates including (D) phenylalanine and tryptophan or (E) leucine and isoleucine. Abbreviations: Phe, phenylalanine; Trp, tryptophan; Leu, leucine; Ile, isoleucine; PAOx, phenylacetaldoxime; IAOx, indole-3-acetaldoxime; 3-MBOx, 3-methylbutanaloxime; 2-MBOx, 2-methylbutanaloxime. Aldoxime formation was monitored by LC-MS analysis. In (D), chromatograms in the bottom rows represent authentic standards used to confirm the identities of PAOx and IAOx. In (E), the chemical identities of 3-MBOx and 2-MBOx were validated by LC-MS/MS fragmentation analysis, as shown in Supplementary Fig. S3.

To determine their catalytic activities, recombinant proteins of the identified SlCYP79s were heterologously expressed in *E. coli* and incubated in auto-induction medium supplemented with each amino acid substrate. Of the various amino acids tested, LC-MS analysis of the reaction products showed that SlCYP79DB32 and SlCYP79DB52 converted phenylalanine into PAOx, whereas only SlCYP79DB52 converted tryptophan into IAOx (Fig. 1D). When tested with aliphatic amino acids, SlCYP79DB52 converted both leucine and isoleucine into 3-methylbutanaloxime (3-MBOx) and 2-methylbutanaloxime (2-MBOx), respectively (Fig. 1E; Supplementary Fig. S3). SlCYP79DB32 produced 2-MBOx from isoleucine but showed no detectable activity with leucine (Fig. 1E; Supplementary Fig. S3).

### Aldoxime overproduction alters tomato growth and increases auxin accumulation

SlCYP79DB32 and SlCYP79DB52 showed broad substrate specificity (Fig. 1D and 1E), making it difficult to selectively elevate a specific aldoxime using the endogenous tomato enzymes. To investigate the physiological consequences of aldoxime accumulation in planta, we therefore employed Arabidopsis CYP79A2 and CYP79B2, which are more substrate-specific. AtCYP79A2 converts Phe to PAOx, whereas AtCYP79B2 converts Trp to IAOx ^8,20^. These genes were introduced into tomato using the pOpON inducible system ^38^, in which dexamethasone (dex) activates LhGR, a dex-responsive chimeric transcription factor, resulting in bidirectional transcription activation of the target gene and the GUS marker gene ^39,40^.

Two single-insertion homozygous lines were established for each construct: *pOpON:AtCYP79A2-2* (*pOpA2-2*) and *pOpON:AtCYP79A2-3* (*pOpA2-3*) for *CYP79A2,* and *pOpON:AtCYP79B2-12* (*pOpB2-12*) and *pOpON:AtCYP79B2-15* (*pOpB2-15*) for *CYP79B2*. In *pOpA2-2* and *pOpA2-3,* GUS staining was detected only after dex treatment, whereas no staining was observed in mock-treated transgenic lines or in wild-type plants with or without dex, confirming dex-dependent activation of *AtCYP79A2* (Fig. 2A). Under mock conditions, the *pOpA2* transgenic lines were morphologically similar to wild type (Fig. 2B). Upon dex application, however, both *pOpA2-2 and pOpA2-3* plants showed increased stem height compared with their mock-treated counterparts, while dex treatment had no effect on wild-type plants (Fig. 2B and 2C). Consistent with the known activity of AtCYP79A2, dex-treated *pOpA2* lines accumulated PAOx, which was undetectable in mock-treated controls (Fig. 2D). Because PAOx has been proposed as an intermediate in PAA biosynthesis, we next examined whether PAOx accumulation was associated with increased PAA levels, previously shown to promote shoot elongation when exogenously applied to tomato ^41^. Both *pOpA2-2* and *pOpA2-3* plants showed significantly elevated endogenous PAA concentrations (Fig. 2E). To further test whether PAOx can be converted to PAA in vivo, we fed wild-type tomato seedlings with either unlabeled PAOx or phenyl ring deuterium-labeled PAOx (D_5_-PAOx). PAOx feeding increased endogenous PAA levels relative to water-fed controls (Fig. 2F), and D_5_-PAA was detected in seedlings supplied with D_5_-PAOx (Fig. 2G), supporting PAOx as a precursor for PAA biosynthesis in tomato.

**Figure 2.**
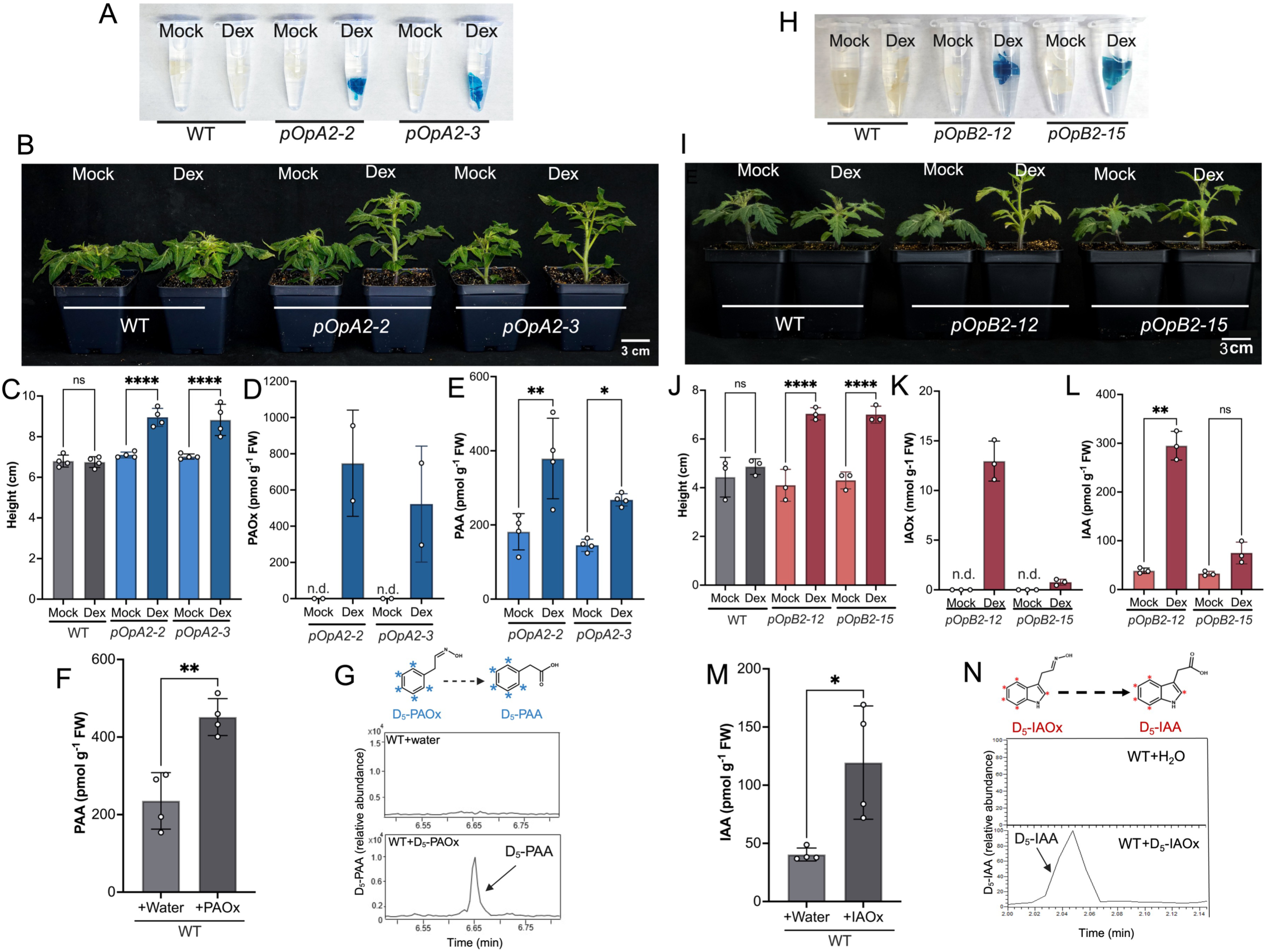
Characterization of inducible aldoxime-overproducing tomato lines and in vivo conversion of aldoximes to auxins. (A–G) Characterization of *pOpA2* lines and PAOx-derived PAA biosynthesis. (A) GUS histochemical staining of cotyledon explants from wild-type and *pOpA2* transgenic lines after treatment with dexamethasone (Dex) or mock solution. Blue staining indicates dex-dependent induction of the pOp/LhGR system. (B) Representative images of 3-week-old wild-type and *pOpA2* plants treated with mock or dex. Scale bar, 3 cm. (C) Quantification of stem height in plants shown in (B). (D, E) PAOx (D) and PAA (E) levels in mock– and dex-treated *pOpA2* leaves. (F) PAA levels in 2-week-old wild-type seedlings fed with water or PAOx for 24 h. (G) Conversion of D_5_-PAOx to D_5_-PAA in wild-type seedlings. The schematic shows D_5_-PAOx and D_5_-PAA structures, with blue asterisks indicating deuterium atoms. Representative GC-MS extracted ion chromatograms show D_5_-PAA detection in D_5_-PAOx-fed seedlings compared with the water control. (H–N) Characterization of *pOpB2* lines and IAOx-derived IAA biosynthesis. (H) GUS histochemical staining of cotyledon explants from wild-type and *pOpB2* transgenic lines after dex or mock treatment. (I) Representative images of 3-week-old wild-type and *pOpB2* plants treated with mock or dex. Scale bar, 3 cm. (J) Quantification of stem height in plants shown in (I). (K, L) IAOx (K) and IAA (L) levels in mock– and dex-treated *pOpB2* leaves. (M) IAA levels in 2-week-old wild-type seedlings fed with water or IAOx for 24 h. (N) Conversion of D_5_-IAOx to D_5_-IAA in wild-type seedlings. The schematic shows D_5_-IAOx and D_5_-IAA structures, with red asterisks indicating deuterium atoms. Representative LC-MS extracted ion chromatograms show D_5_-IAA detection in D_5_-IAOx-fed seedlings compared with the water control. For bar graphs, individual data points are shown, and error bars represent SD; biological replicate numbers are indicated by the plotted data points in each panel. “n.d.” indicates not detected. Asterisks indicate statistically significant differences determined by Student’s t-test (*P < 0.05, **P < 0.01, ****P < 0.0001; ns, not significant).

We next examined the Trp-derived IAOx-to-IAA pathway using *pOpB2* lines. In *pOpB2-12* and *pOpB2-15* plants, GUS staining was induced by dex treatment, confirming dex-specific activation of the *AtCYP79B2* transgene (Fig. 2H). Dex-treated *pOpB2* plants also displayed increased stem height compared with mock-treated controls, whereas wild-type plants showed no height response to dex treatment (Fig. 2I and 2J). This phenotype is reminiscent of previously reported auxin-overproduction phenotypes in Arabidopsis *CYP79B2*-overexpressing plants, which showed elongated hypocotyls and elevated free IAA levels ^42^. Metabolite profiling showed that dex-treated *pOpB2* lines accumulated IAOx, which was undetectable in mock-treated controls, together with significantly elevated IAA concentrations (Fig. 2K and 2L). To determine whether this aldoxime intermediate can contribute to IAA biosynthesis, we fed wild-type seedlings with exogenous IAOx. IAOx feeding increased endogenous IAA levels (Fig. 2M), and isotope-tracing experiments detected the conversion of D_5_-IAOx to D_5_-IAA (Fig. 2N), demonstrating that IAOx can serve as a precursor of IAA in tomato. Together, these results show that PAOx and IAOx can be converted to their corresponding auxins, PAA and IAA, in tomato, and that induced aldoxime accumulation is associated with increased auxin accumulation.

### SlNIT1 catalyzes the conversion of benzyl cyanide to PAA

While the enzymatic steps mediating the IAOx-to-IAA conversion remain unclear ^43^, several studies suggest that benzyl cyanide (BC) may serve as an intermediate of the PAOx-PAA pathway ^9,44^. Consistent with this, we found substantial accumulation of BC in the dex-induced *pOpA2* lines compared to mock-treated plants (Fig. 3A). GC-MS analysis detected D_5_-BC in tomato seedlings fed with D_5_-PAOx, while no D_5_-BC signal was detected in water-fed controls, indicating that BC is derived from PAOx in vivo (Fig. 3B). To test whether BC is subsequently converted to PAA, we fed tomato seedlings with ^13^C-BC, labeled with ^13^C at the α-carbon. ^13^C-PAA was detected in ^13^C-BC-fed seedlings but not in water-fed controls (Fig. 3C), confirming that BC serves as an intermediate in the PAOx-derived PAA biosynthetic pathway in tomato.

**Figure 3.**
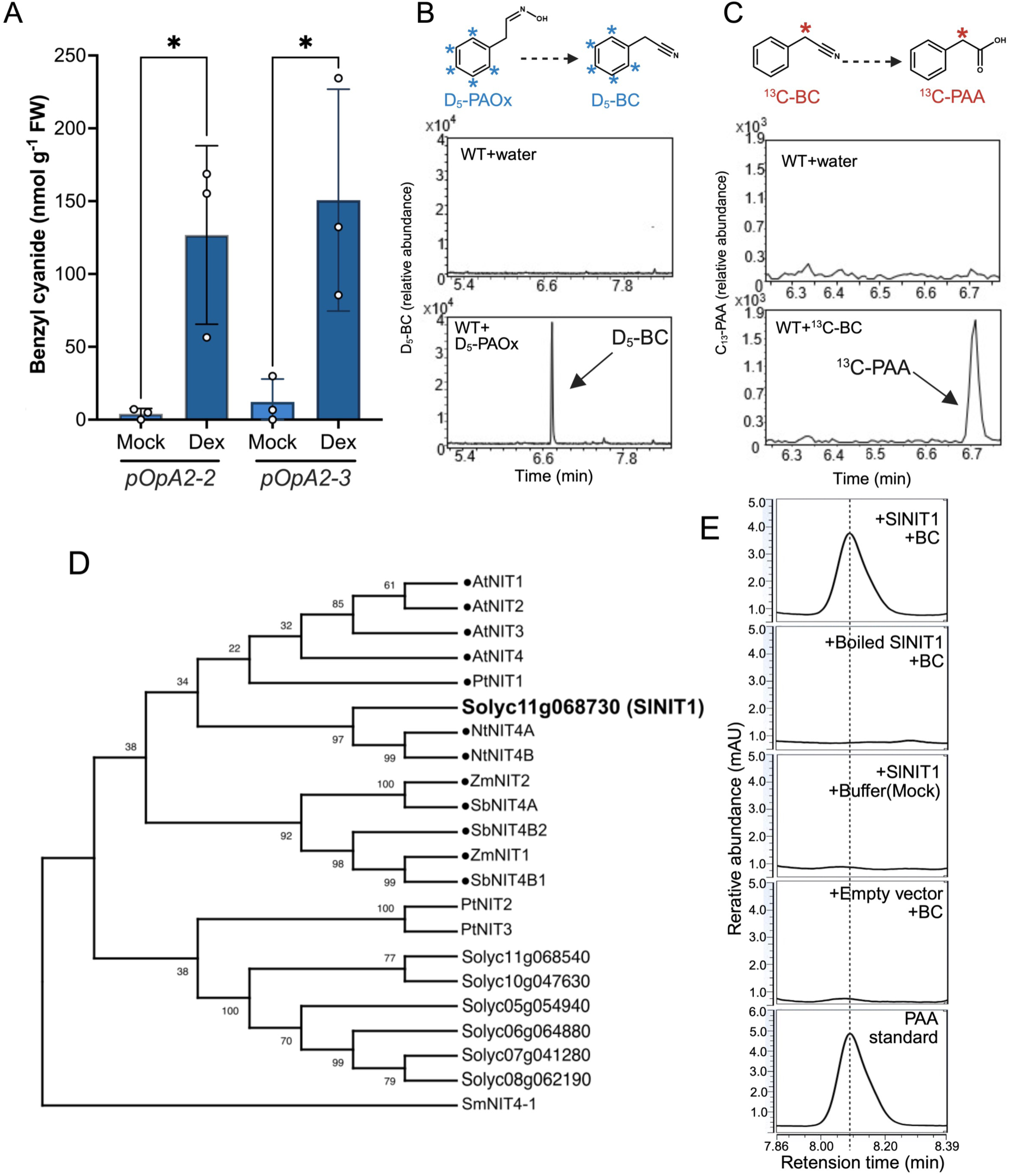
Identification of benzyl cyanide (BC) as an intermediate in the PAOx to PAA pathway in tomato and functional characterization of SlNIT1. (A) Accumulation of BC in tomato *pOpA2* lines. BC content was measured in the leaves of the inducible *pOpA2* lines following treatment with mock or dexamethasone (Dex) solutions. Data represent the mean ± SD (n = 3) and asterisks indicate statistically significant differences determined by Student’s t-test (*P<0.05). (B) In vivo tracing of the PAOx to BC conversion. The schematic illustrates the structure of D_5_-PAOx and the expected D_5_-BC product where blue asterisks indicate deuterium atoms. Representative GC-MS extracted ion chromatograms demonstrate the accumulation of D_5_-BC in two-week-old wild-type seedlings fed with D_5_-PAOx for 24 hours compared to the water-fed control shown in the top row. (C) In vivo conversion of BC to PAA. The schematic shows the conversion of ^13^C-labeled BC to ^13^C-labeled PAA where the red asterisk indicates the ^13^C label. Representative GC-MS extracted ion chromatograms at m/z 152 show specific detection of ^13^C-PAA in two-week-old wild-type seedlings fed ^13^C-BC for 24 hours, compared with the water control shown in the top row. (D) Phylogenetic analysis of plant nitrilases. The Maximum Likelihood tree with 500 bootstrap replicates includes seven identified tomato homologs (SlNITs) and characterized NIT proteins from *Arabidopsis thaliana* (At), *Zea mays* (Zm), *Sorghum bicolor* (Sb), *Nicotiana tabacum* (Nt), and *Populus trichocarpa* (Pt). *Selaginella moellendorffii* (Sm) NIT4-1 serves as the outgroup. (E) In vitro enzymatic activity of SlNIT1. Recombinant SlNIT1 was expressed in *E. coli* and the crude extracts were incubated with BC. HPLC chromatograms displaying UV absorbance in mAU show the production of PAA in the SlNIT1 reaction, matching the authentic PAA standard in the bottom row. Controls included boiled enzyme extracts, buffer-only, and empty vector extracts.

Nitrilases catalyze the hydrolysis of nitriles to their corresponding carboxylic acids ^44–47^, making them candidate enzymes for the conversion of BC to PAA. We identified seven putative nitrilase coding genes in the tomato genome and conducted phylogenetic analysis together with previously characterized nitrilases from other plant species (Fig. 3D). Among the seven candidates, Solyc11g068730 clustered with functionally validated nitrilases and shared substantial sequence similarity (>60% identity) with characterized nitrilases, while the remaining six candidates exhibited less than 30% sequence identity (Fig. 3D; Supplementary Fig. S4). Based on these analyses, we designated Solyc11g068730 as SlNIT1. To determine whether SlNIT1 can hydrolyze BC, we expressed recombinant SlNIT1 in *E. coli* and incubated it with this substrate. HPLC analysis revealed a distinct peak corresponding to PAA in the SlNIT1 reaction with BC, whereas no PAA was detected in the controls, including boiled enzyme, buffer-only, and empty-vector extracts (Fig. 3E). These results demonstrate that SlNIT1 catalyzes the conversion of BC to PAA in vitro, supporting its possible role in the PAOx-derived PAA biosynthesis in tomato.

### Aldoxime overproduction reduces phenylpropanoid accumulation in tomato

During the cultivation of the inducible transgenic lines, we observed that dex-treated *pOpA2* and *pOpB2* plants exhibited markedly reduced hypocotyl pigmentation compared with mock controls (Fig. 4A and 4B). Because this pigmentation is typically attributed to anthocyanin accumulation, we quantified total anthocyanin levels. Spectrophotometric analysis confirmed a significant reduction in anthocyanin content in dex-treated *pOpA2* and *pOpB2* lines relative to their respective mock controls, while wild-type plants showed no difference upon dex application (Fig. 4C-F).

**Figure 4.**
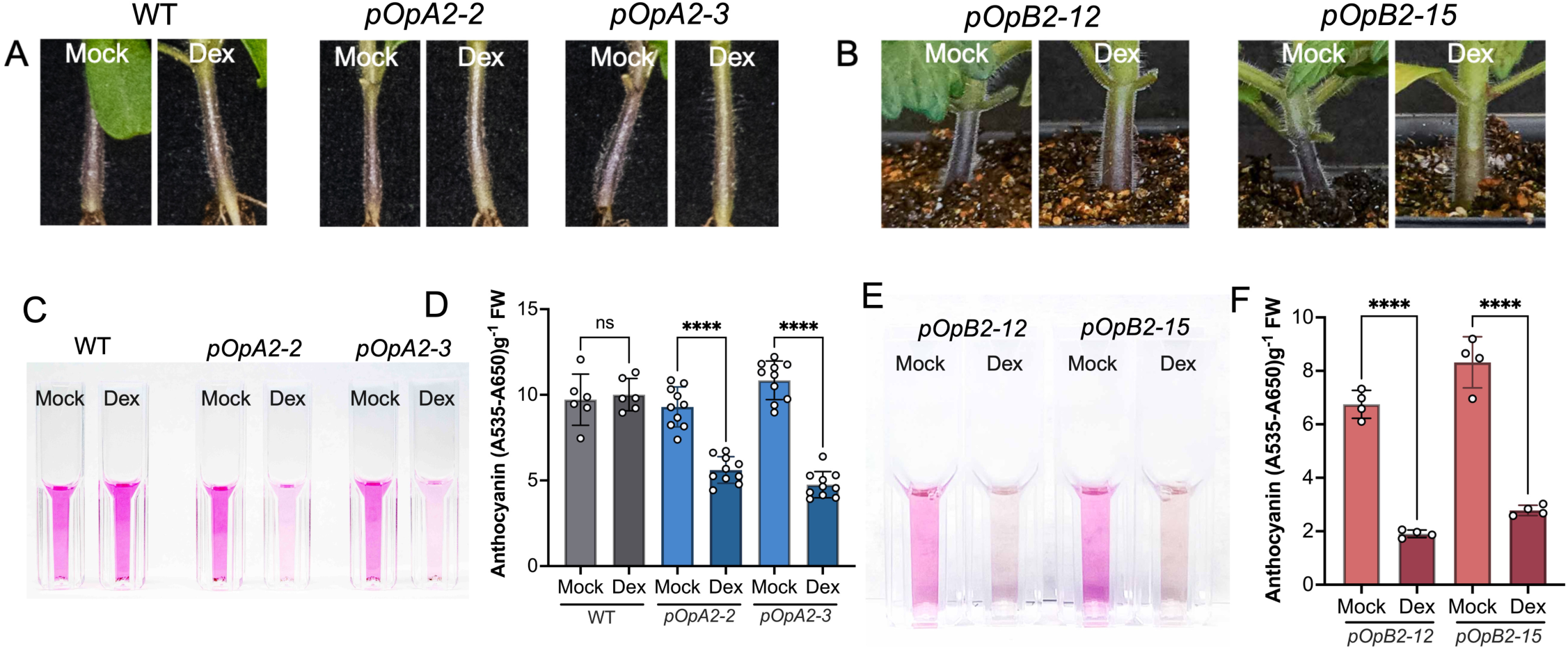
Reduced anthocyanin accumulation in aldoxime-overproducing tomato. (A, B) Representative close-up images of the basal stem region of (A) *pOpA2* and (B) *pOpB2* transgenic lines compared to wild-type plants. Plants were treated with a mock solution or dexamethasone (Dex) to induce transgene expression. Under mock conditions, stems of both transgenic and wild-type plants display typical purple pigmentation indicative of anthocyanin accumulation. In contrast, dex-treated *pOpA2* and *pOpB2* plants exhibit a distinct loss of purple coloration. (C, E) Visual comparison of anthocyanin extracts from two-week-old soil-grown seedlings of (C) *pOpA2* and (E) *pOpB2* transgenic lines. Plants were treated with a mock solution or 20 µM dex sprayed twice over six days. The loss of pink pigmentation in dex-treated samples indicates a reduction in anthocyanin content. (D, F) Spectrophotometric quantification of total anthocyanin levels, expressed as A535−A650 g^-1^ FW, in (D) *pOpA2* and (F) *pOpB2* lines. Data represent the mean ± SD where n = 10 for *pOpA2*, n = 4 for *pOpB2*, and n = 6 for wild-type plants. Asterisks indicate statistically significant differences determined by Student’s t-test (****P < 0.0001; ns, not significant).

To investigate if other phenylpropanoids were affected, we profiled soluble phenylpropanoids in methanol extracts from dex-treated and mock-treated *pOpA2* and *pOpB2* leaves by HPLC. Chromatograms revealed an overall reduction in peak intensity across multiple peaks in the dex-treated samples of both lines (Fig. 5A and 5B). Using authentic standards, we identified two of the prominently reduced metabolites as chlorogenic acid and rutin. Quantitative analysis confirmed that both chlorogenic acid and rutin levels were significantly decreased in dex-treated *pOpA2* and *pOpB2* lines compared to mock controls (Fig. 5C-F). These results indicate that aldoxime overproduction reduces phenylpropanoid accumulation in tomato.

**Figure 5.**
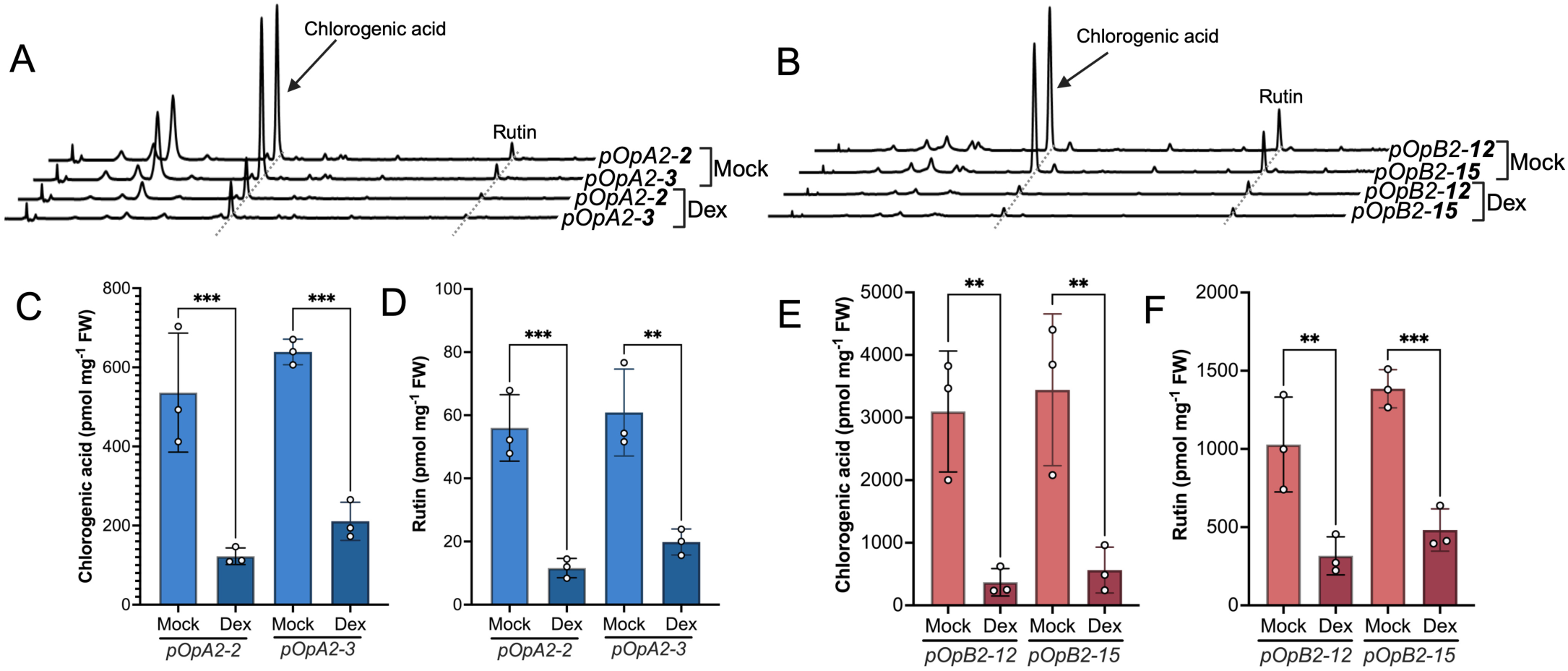
Suppression of soluble phenylpropanoid accumulation in aldoxime-overproducing tomato. (A, B) Representative HPLC chromatograms of soluble phenylpropanoids in (A) *pOpA2* and (B) *pOpB2* samples. For *pOpA2* lines, seedlings grown on half-strength MS medium were treated with 20 μM dexamethasone (Dex) or mock for two weeks. For *pOpB2* lines, leaves were collected from three-week-old soil-grown plants after seven days of dex or mock treatment. Peaks corresponding to chlorogenic acid and rutin are indicated and show a marked reduction under dex-induced conditions compared with mock controls. (C, D) Quantitative analysis of (C) chlorogenic acid and (D) rutin levels in *pOpA2* lines. (E, F) Quantitative analysis of (E) chlorogenic acid and (F) rutin levels in *pOpB2* lines. Data represent the mean ± SD of three biological replicates (n = 3). Asterisks indicate statistically significant differences determined by Student’s t-test (**P < 0.01, ***P < 0.001).

Because Phe serves as the entry-point substrate for the phenylpropanoid pathway (Fig. 6A) and as the precursor for PAOx biosynthesis, it is possible that competitive depletion of the shared precursor accounted for the reduced phenylpropanoid levels in the *pOpA2* lines. However, supplementation with exogenous Phe failed to restore chlorogenic acid or rutin levels in dex-treated *pOpA2* seedlings (Supplementary Fig. S5). In Arabidopsis, aldoxime-mediated phenylpropanoid repression is primarily regulated through the targeted degradation of phenylalanine ammonia-lyase (PAL), mediated by transcriptional activation of Kelch repeat F-box (KFB) proteins ^24,25^. To determine whether a similar regulatory mechanism operates in tomato, we measured total PAL enzymatic activity. Unexpectedly, PAL activity was not reduced in dex-treated *pOpA2* and *pOpB2* lines. Instead, it was either elevated or unchanged (Fig. 6B and 6C), indicating that the suppression of phenylpropanoid accumulation is not attributable to reduced PAL activity.

**Figure 6.**
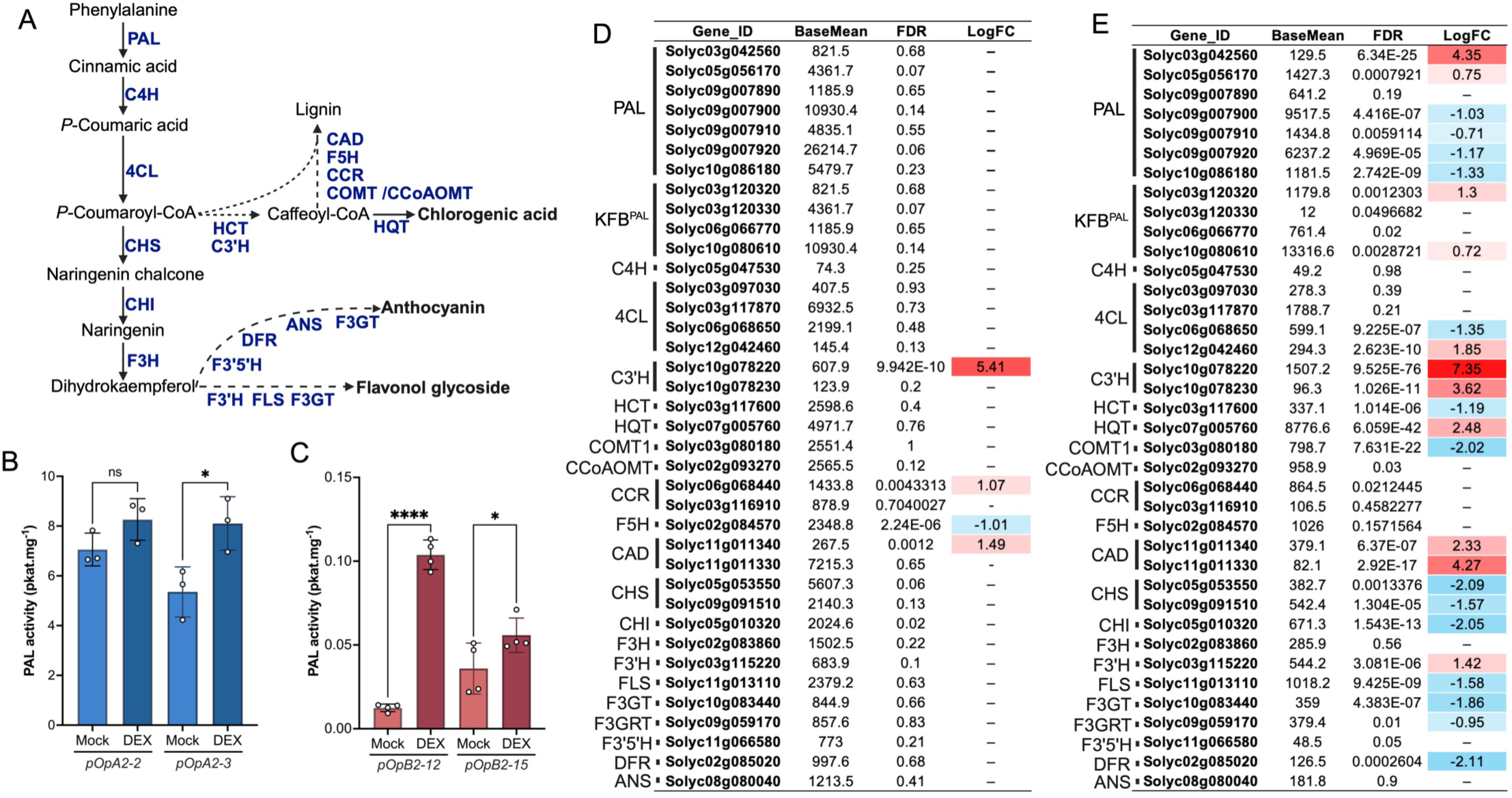
Effect of aldoxime overproduction on the phenylpropanoid pathway in tomato. (A) Schematic representation of the general phenylpropanoid pathway in tomato. Key enzymes are abbreviated as follows. PAL, phenylalanine ammonia-lyase. C4H, cinnamate 4-hydroxylase. 4CL, 4-coumarate-CoA ligase. HCT, hydroxycinnamoyl-CoA shikimate/quinate hydroxycinnamoyltransferase. C3’H, p-coumaroyl ester 3’-hydroxylase. CHS, chalcone synthase. CHI, chalcone isomerase. F3H, flavanone 3-hydroxylase. F3’H, flavonoid 3’-hydroxylase. F3’5’H, flavonoid 3’,5’-hydroxylase. FLS, flavonol synthase. DFR, dihydroflavonol 4-reductase. ANS, anthocyanidin synthase. F3GT, flavonoid 3-O-glucosyltransferase. CAD, cinnamyl alcohol dehydrogenase. F5H, ferulate 5-hydroxylase. CCR, cinnamoyl-CoA reductase. COMT, caffeic acid O-methyltransferase. CCoAOMT, caffeoyl-CoA O-methyltransferase. HQT, hydroxycinnamoyl-CoA quinate hydroxycinnamoyltransferase. (B) PAL enzymatic activity in *pOpA2* lines (*pOpA2-2* and *pOpA2-3*). Seedlings were grown in liquid half-strength MS medium for one week, followed by treatment with 20 μM dex or mock solution for one week. (C) PAL enzymatic activity in *pOpB2* lines (*pOpB2-12* and *pOpB2-15*). Total protein was extracted from the leaves of three-week-old soil-grown plants treated with dex spray or mock for one week. (D) Transcriptional profiling table of PAL, KFB homologs targeting PAL (KFB^PAL^), and structural genes of the general phenylpropanoid pathway in *pOpA2* lines. (E) Transcriptional profiling table of PAL, KFB^PAL^ homologs, and structural genes of the general phenylpropanoid pathway in *pOpB2* lines. For the transcriptomic data tables (D and E), values display BaseMean, false discovery rate (FDR), and log_2_ fold change (LogFC) for Dex versus Mock. Highlighted cells in the LogFC column indicate significant differential expression at FDR < 0.01. Genes with FDR ≥ 0.01 are indicated by a dash (–) in the LogFC column. Data in bar graphs represent the mean ± SD (n = 3). Asterisks indicate statistically significant differences determined by Student’s t-test where * indicates P<0.05, **** indicates P<0.0001, and ns denotes not significant.

To further explore the basis of this suppression, we examined the transcriptional profiles of phenylpropanoid pathway genes using RNA-seq. In the *pOpA2* lines, PAL and KFB^PAL^ homologs did not show significant changes between mock and dex treatments using an FDR cutoff of 0.01, and most phenylpropanoid biosynthetic genes were unchanged (Fig. 6D). In the *pOpB2* lines, PAL and KFB^PAL^ homologs showed mixed responses, with two PAL homologs upregulated, four downregulated, and one unchanged. Among the KFB^PAL^ homologs, two of four were upregulated, whereas the remaining two were unchanged (Fig. 6E). These transcriptional patterns did not provide a simple explanation for the broad reduction in phenylpropanoid metabolites observed in the aldoxime-overproducing lines (Fig. 6D and 6E). As the transcriptional data do not reveal a clear or consistent regulatory pattern that would account for the observed suppression of phenylpropanoid accumulation, it is possible that additional regulatory layers, such as post-transcriptional control or alternative metabolic regulatory mechanisms, may be involved.

### Transcriptome analysis reveals altered expression of stress-related genes

Beyond the phenylpropanoid pathway, we next examined the broader transcriptional landscape to gain a comprehensive view of the global changes induced by aldoxime overproduction. First, we examined DEGs associated with auxin biology to corroborate the physiological and metabolic evidence for enhanced auxin activity. Consistent with the elevated auxin levels and increased stem elongation observed in the transgenic lines (Fig. 2), genes involved in auxin response, signaling, and transport were transcriptionally activated in both *pOpA2* and *pOpB2* lines (Supplementary Fig. S6). This upregulation was substantially broader in the *pOpB2* lines, where a larger number of auxin-responsive, Aux/IAA, ARF, and PIN/LAX family genes were induced compared to *pOpA2* (Supplementary Fig. S6). This difference in transcriptional breadth may reflect the distinct biological activities and metabolic contexts of IAA and PAA.

To identify the biological processes affected, we performed Gene Ontology (GO) analyses, which revealed that both *pOpA2* and *pOpB2* lines showed significant enrichment of terms associated with stress and defense responses, including “defense response” and “response to stress” (Fig. 7A and 7B). Given that aldoximes and their derivatives are related to defensive and stress responses in other plant species ^48,10,36^, it is possible that aldoxime metabolism may function in stress adaptation in tomato. Downregulated DEGs in both lines were predominantly enriched in primary metabolism categories, including “photosynthesis,” “chloroplast organization,” and “carbon metabolism” (Fig. 7C and 7D). This widespread transcriptional suppression of primary metabolic processes suggests a global reallocation of resources from growth-related metabolism toward defense and specialized metabolism upon aldoxime overproduction.

**Figure 7.**
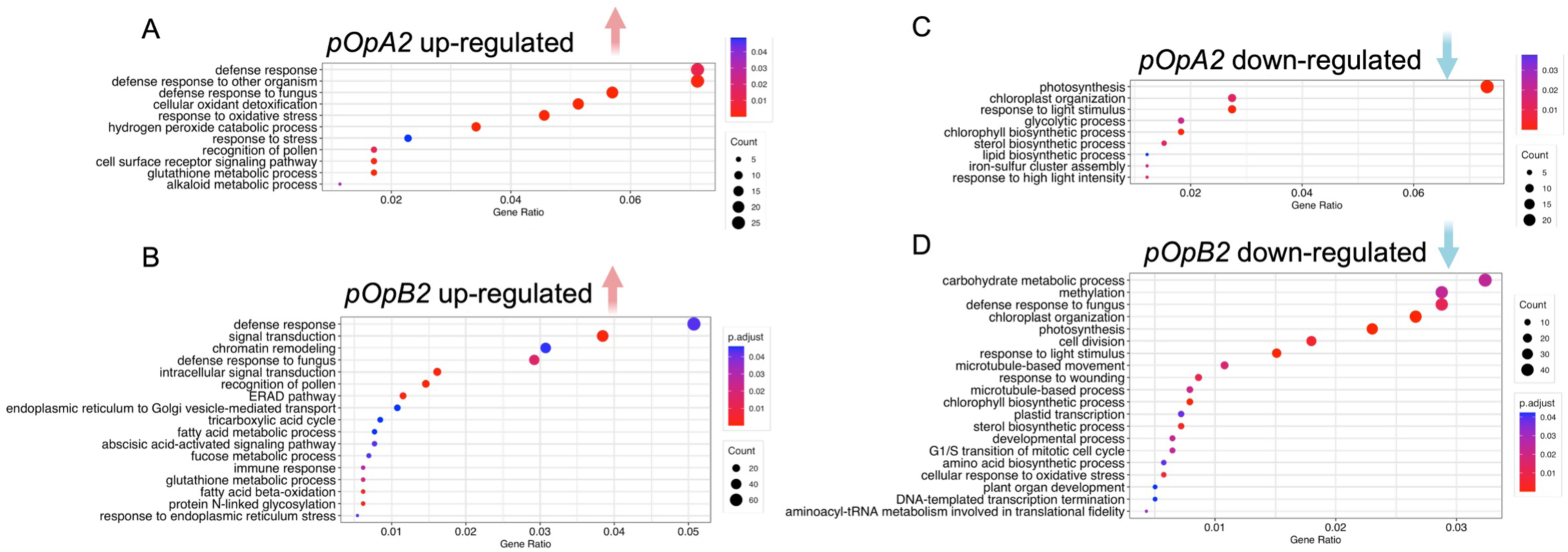
Transcriptional reprogramming and metabolic shifts in aldoxime-overproducing tomato lines. (A–D) Functional enrichment analysis of differentially expressed genes. Bubble plots display the top enriched Gene Ontology biological processes upregulated in (A) *pOpA2* and (B) *pOpB2*, and downregulated in (C) *pOpA2* and (D) *pOpB2* lines upon dexamethasone induction. Differentially expressed genes were defined using an adjusted P-value cutoff of 0.05. The color gradient from red to blue represents the adjusted P-value (p.adjust) calculated via the Benjamini-Hochberg correction, and the dot size indicates the number of genes (Count) mapped to each term. The enrichment pattern indicates that stress– and defense-related terms are enriched among upregulated genes, whereas photosynthesis and primary metabolism terms are predominantly enriched among downregulated genes.

## Discussion

Aldoximes represent important branch points connecting amino acid metabolism with growth hormone auxins and specialized metabolism, yet their roles remain poorly understood in many plant species. Here, using tomato as a model solanaceous crop, we provide biochemical and genetic evidence that aldoxime metabolism contributes to auxin biosynthesis and modulates phenylpropanoid accumulation, revealing a previously uncharacterized metabolic connection in this crop species.

Five putative CYP79-encoding genes were identified in the tomato genome. Among them, SlCYP79DB52 showed broad substrate specificity toward both aromatic (Phe, Trp) and aliphatic (Leu, Ile) amino acids, whereas SlCYP79DB32 displayed partial selectivity, accepting Phe and Ile but not Trp or Leu (Fig. 1D and 1E). This biochemical differentiation, together with distinct spatiotemporal expression patterns observed in available transcriptomic datasets (Fig. 1C), suggests potential non-redundant roles in tomato metabolism. The remaining three CYP79 homologs (SlCYP79DB43, SlCYP79DB46, SlCYP79DB47) showed no detectable activity toward the tested substrates under our experimental setting. These enzymes may require untested substrates, specific reaction conditions, or protein partners for activity, although the possibility that some represent nonfunctional or weakly active homologs cannot be excluded. More broadly, CYP79 enzymes exhibit varying degrees of substrate plasticity across plant lineages. For example, several CYP79 enzymes from poplar and neotropical myrmecophyte tococa exhibit broad substrate specificity ^10,36^, while Arabidopsis AtCYP79A2 and AtCYP79B2/B3 show strict specificity for Phe and Trp, respectively ^7,8^. It is noteworthy that CYP79 enzymes are absent in certain plant lineages, such as ferns and orchids, where flavin-containing monooxygenases catalyze aldoxime formation ^4,49^. Collectively, these findings suggest that plant lineages have evolved distinct enzymatic strategies for aldoxime biosynthesis, differing in both the enzyme families employed and their substrate specificities.

Our in vivo feeding and isotope-tracing results demonstrate that IAOx and PAOx can be converted into IAA and PAA, respectively, in tomato (Fig. 2). For the IAOx-to-IAA conversion, the biosynthetic route remains unresolved. Earlier isotope-labeling studies in Arabidopsis detected labeled indole-3-acetonitrile (IAN) and indole-3-acetamide (IAM), suggesting that these compounds may act as intermediates connecting IAOx to IAA ^21^. However, IAN can also arise from the turnover of indole glucosinolates, complicating the interpretation of this route ^50^. More recent genetic evidence indicates that several candidate gene families, including CYP71A, NIT, AMI, and IAMH, are dispensable for IAOx-mediated IAA overproduction in the Arabidopsis *sur2* background, suggesting that additional uncharacterized routes may exist ^43^. Importantly, since tomato lacks the glucosinolate pathway, the metabolic context for IAOx metabolism may differ from that in Brassicaceae, and the enzymes responsible for IAOx-to-IAA conversion in tomato remain unknown.

In contrast, the PAOx-to-PAA route appears to involve a nitrile intermediate. Our data show that PAOx can be converted to benzyl cyanide (BC), which can subsequently be hydrolyzed to PAA (Fig. 3A-C). The identification and functional validation of SlNIT1 as a nitrilase capable of hydrolyzing BC to PAA supports the presence of a nitrilase-dependent step in the PAOx-derived PAA biosynthetic route in tomato (Fig. 3D and 3E). However, how BC is synthesized from PAOx remains unknown. Several studies have suggested that CYP71 enzymes catalyze the conversion of aldoximes to their corresponding nitriles in multiple plant species ^36,51,52^. Consequently, tomato CYP71 enzymes might be involved in the formation of BC from PAOx. Alternative routes for BC biosynthesis have been proposed in tomato. Previous genetic evidence suggested a phenethylamine-dependent pathway, as manipulation of tomato aromatic amino acid decarboxylase genes altered BC levels ^53^. More recently, isotope-labeling experiments in tomato fruits suggested that BC can also be synthesized via cysteine-derived thiazolidine intermediates, with cysteine serving as the nitrogen source rather than Phe^54^. Our study demonstrates that BC can be produced from Phe via PAOx in tomato leaves, suggesting that tomato may use multiple tissue– or developmental context-dependent routes for BC biosynthesis. Further work is needed to determine the relative contributions of these pathways to BC and PAA production.

Notably, PAOx and IAOx were not detected in mock-treated tomato leaves under our standard growth conditions (Fig. 2D and 2K). This is consistent with the low expression of *SlCYP79DB52* and *SlCYP79DB32* in leaves, suggesting that leaf tissue under non-stressed conditions may not be a major site of aldoxime accumulation. A similar pattern has been reported for *AtCYP79A2* in Arabidopsis, where low expression across most organs under optimal conditions is associated with the absence of benzyl glucosinolate accumulation in the Col-0 ecotype ^22,55^. Indeed, in many plant species, aldoxime production is not constitutive but is instead induced by environmental stimuli, with *CYP79* genes frequently upregulated in response to stresses such as high temperature, herbivory, and jasmonic acid treatment ^10,35,36,56^. For example, *ZmCYP79A61* expression is strongly induced by herbivore attack in maize, accompanied by elevated levels of PAOx and PAA ^57^. Taken together, these observations suggest that aldoxime metabolism in tomato is likely restricted to specific tissues, developmental stages, or stress conditions rather than being broadly active under standard growth conditions. Future studies across diverse environmental conditions, developmental stages, and tomato genetic backgrounds beyond Micro-Tom will help define the physiological roles of aldoxime metabolism in tomato.

Among the findings of this study, aldoxime overproduction led to a pronounced suppression of phenylpropanoid accumulation (Figs. 4 and 5). Because PAOx is derived from Phe, one possible explanation is that PAOx overproduction diverts Phe away from phenylpropanoid biosynthesis. However, exogenous Phe supplementation failed to rescue the reduced phenylpropanoid accumulation in the *pOpA2* lines (Supplementary Fig. S5). Phenylpropanoid suppression was also observed in *pOpB2* lines, where the overexpressed *CYP79B2* uses Trp rather than Phe as its substrate. These results argue against simple precursor competition as the primary mechanism underlying phenylpropanoid suppression. In Arabidopsis, aldoxime-mediated phenylpropanoid suppression has been linked to transcriptional upregulation of KFB proteins, which mediate ubiquitin-dependent proteasomal degradation of PAL and thereby reduce metabolic flux into the phenylpropanoid pathway ^24,25^. Our results in tomato do not fit this model. PAL enzymatic activity was elevated or unchanged, rather than reduced, in dex-treated *pOpA2* and *pOpB2* lines (Fig. 6B and 6C). Furthermore, transcriptomic analysis did not reveal a consistent pattern of KFB upregulation, PAL downregulation, or coordinated repression of downstream phenylpropanoid biosynthetic genes that could fully account for the broad reduction in phenylpropanoid accumulation (Fig. 6D and 6E). Together, these observations suggest that aldoxime accumulation in tomato suppresses phenylpropanoid production through mechanisms distinct from simple substrate depletion or the canonical Arabidopsis KFB-PAL model. The suppression may involve regulatory steps downstream of PAL in the phenylpropanoid pathway, post-transcriptional or post-translational control, or broader metabolic regulation.

RNA-seq analysis further revealed transcriptional reprogramming involving auxin-and defense-associated pathways upon aldoxime overproduction (Fig. 7; Supplementary Fig. S6). Consistent with the increased auxin accumulation and stem elongation observed in the transgenic lines, genes involved in auxin response, signaling, and transport were transcriptionally activated, particularly in *pOpB2* lines (Supplementary Fig. S6). Beyond these auxin-related changes, the upregulation of stress– and defense-related genes, together with the downregulation of photosynthesis-and carbon metabolism-related genes, suggests that aldoxime accumulation may shift tomato leaves toward a defense-associated metabolic state (Fig. 7). This pattern is consistent with the established roles of aldoximes as precursors of various defense compounds, including glucosinolates, camalexin, and cyanogenic glucosides, in diverse plant species ^2,33^.

In conclusion, this study identifies aldoxime metabolism as a previously uncharacterized metabolic hub in tomato that links auxin biosynthesis to phenylpropanoid regulation (Fig. 8). By demonstrating that PAOx and IAOx can contribute to PAA and IAA biosynthesis, respectively, our findings expand the known metabolic routes for auxin production in a solanaceous crop. The suppression of phenylpropanoid accumulation further reveals a regulatory connection between aldoxime metabolism and specialized metabolism. Together, these results extend the functional significance of CYP79-dependent aldoxime metabolism beyond Brassicaceae and provide new insight into metabolic networks that coordinate growth-related hormones, defense-associated metabolites, and crop-relevant specialized metabolism in tomato.

**Figure 8.**
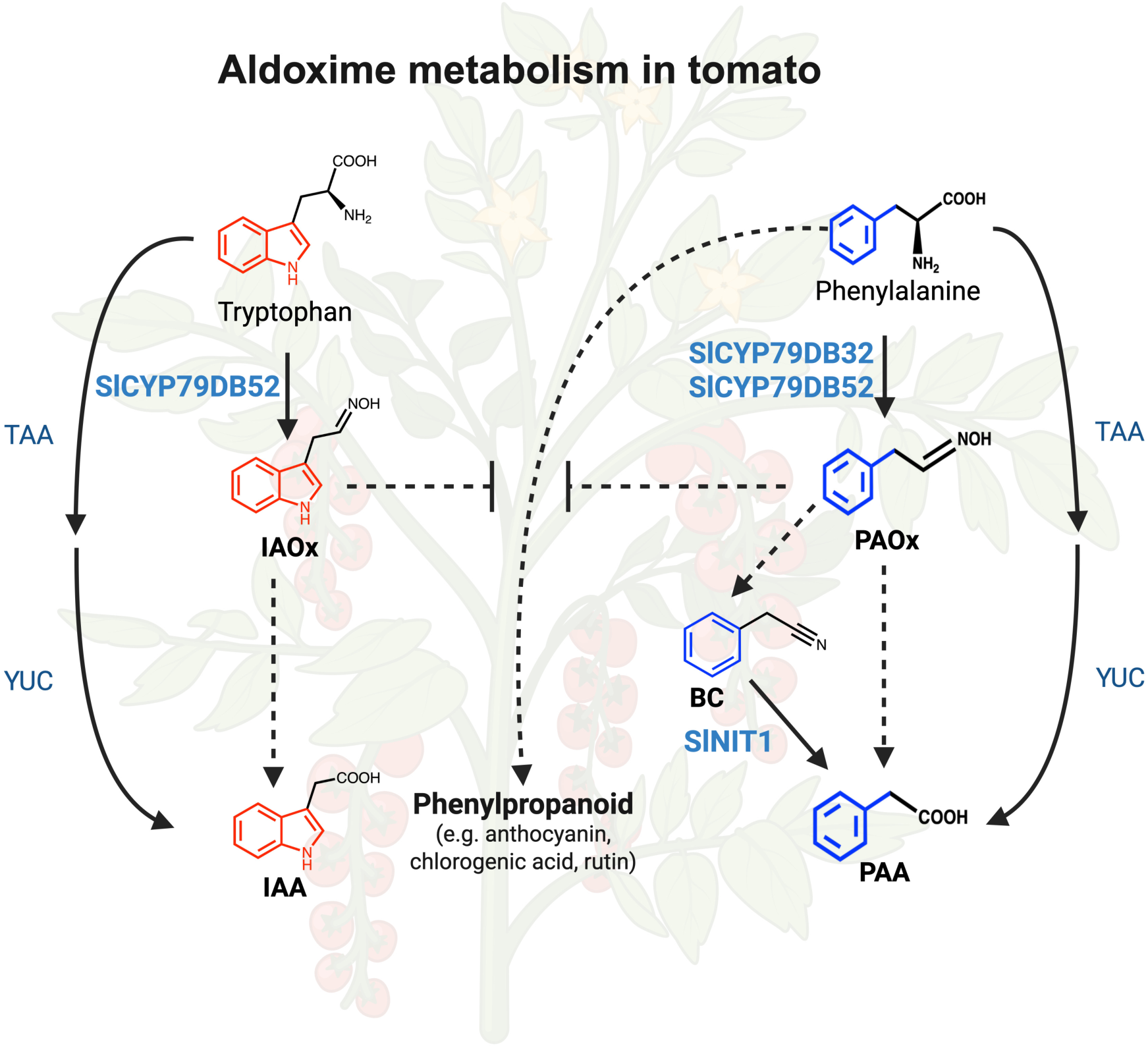
Proposed model for aldoxime-mediated auxin biosynthesis and phenylpropanoid suppression in tomato. Schematic illustration summarizing the metabolic reprogramming observed in this study. SlCYP79DB52 converts tryptophan into indole-3-acetaldoxime (IAOx), whereas SlCYP79DB32 and SlCYP79DB52 convert phenylalanine into phenylacetaldoxime (PAOx). These aldoximes can contribute to the biosynthesis of the auxins indole-3-acetic acid (IAA) and phenylacetic acid (PAA). The canonical TAA/YUC-dependent auxin biosynthetic pathways are shown for comparison. PAOx is converted to PAA via the intermediate benzyl cyanide (BC), and SlNIT1 catalyzes the BC-to-PAA conversion. The accumulation of IAOx and PAOx suppresses the general phenylpropanoid pathway, leading to reduced levels of downstream metabolites, including anthocyanins, chlorogenic acid, and rutin. Solid arrows denote established enzymatic steps, while dashed arrows represent multi-step conversions or regulatory interactions. Enzymes characterized in this study are highlighted in blue.

## Materials and Methods

### Plant Materials and Growth Conditions

Tomato (*Solanum lycopersicum* cv. Micro-Tom) seeds were obtained from the C. M. Rick Tomato Genetics Resource Center (University of California, Davis, USA). Plants were grown at 22 °C under a 16 h light/8 h dark photoperiod. For plate-grown seedlings, seeds were surface-sterilized in 20% (v/v) bleach for 15 min and rinsed several times with sterile water. Sterilized seeds were sown on half-strength Murashige and Skoog (MS) medium supplemented with 2% (w/v) sucrose and 0.8% (w/v) agar. For soil-grown plants, seeds were directly sown in potting soil.

### Identification and Phylogenetic Analysis of Tomato CYP79 and Nitrilase Genes

Homologs of nitrilases and cytochrome P450 enzymes in tomato (*Solanum lycopersicum*, taxon ID: 4081) were identified by querying the UniProt protein database using the keywords “nitrilase” and “cytochrome P450.” Retrieved protein sequences were aligned together with reference sequences using the MUSCLE algorithm. Phylogenetic analysis of nitrilase proteins was performed using the maximum likelihood method with the Whelan and Goldman (WAG) model and a gamma distribution (+G4), with 500 bootstrap replicates implemented in MEGA11. For cytochrome P450 protein sequences, the best-fit substitution model (VT+F+G4) was selected using ModelFinder, and phylogenetic trees were constructed using the maximum likelihood method with 1000 bootstrap replicates in IQ-TREE ^58,59^.

### Heterologous Expression and Enzyme Activity Assay of SlCYP79s and SlNIT1

*SlCYP79* coding sequences were cloned into the pETDuet-1 vector containing *Arabidopsis thaliana ATR1* and expressed in *E.coli* ArcticExpress (DE3) RIL cells. Recombinant constructs were verified by sequencing prior to expression. For SlCYP79 enzyme activity assays, cultures were grown using an auto-induction system. Briefly, overnight cultures were inoculated into auto-induction medium containing tryptone (10 g L^-1^), yeast extract (5 g L^-1^), NaCl (5 g L^-1^), glycerol (0.5% (v/v)), glucose (0.05% (w/v)), lactose (0.2% (w/v)), ammonium sulfate (25 mM), MgSO_4_ (1 mM), potassium phosphate buffer (100 mM, KH_2_PO_4_/Na_2_HPO_4_), and trace metal mix, supplemented with ampicillin (100 µg mL^-1^), gentamicin (20 µg mL^-1^), and 0.5 mM 5-aminolevulinic acid. Amino acid substrates were supplied at 10 mM for phenylalanine, tryptophan, leucine, and isoleucine. Cultures were incubated sequentially at 26 °C for 3 h, 12 °C for 48 h, and 16 °C for 48 h with shaking. Whole cultures (cells and medium) were sonicated on ice and centrifuged at 4,000 × g for 10 min at 4 °C. Supernatants were extracted three times with ethyl acetate, and pooled organic phases were dried and resuspended in 50% methanol for metabolite analysis.

The coding sequence of *SlNIT1* was cloned into the pET-28a (+) vector and expressed in *E. coli* ArcticExpress (DE3) RIL cells. Cells were grown in 100 mL auto-induction medium supplemented with kanamycin (50 µg mL⁻¹) and gentamicin (20 µg mL⁻¹) at 16 °C for 48 h, then harvested by centrifugation. The cell pellet was washed with 20 mL of extraction buffer, centrifuged again, and resuspended in 5 mL of extraction buffer. Cells were lysed by sonication on ice (three cycles), and cell debris was removed by centrifugation at maximum speed for 10 min. The resulting supernatant was used as a crude enzyme extract. For enzyme assays, 500 µL of crude extract was incubated with benzyl cyanide (final concentration 1 mM). Reactions were carried out at 30 °C for 2 h with shaking (180 rpm). Reactions were terminated by the addition of 10 µL formic acid, followed by 250 µL acetonitrile. Samples were centrifuged at maximum speed for 10 min, and the supernatant was analyzed by HPLC.

### Chemical Feeding Assay

The feeding assay was performed as described previously ^22^. Two-week-old Micro-Tom seedlings grown aseptically on half-strength MS plates were harvested and submerged in solutions containing 0.005% (v/v) Triton X-100 with or without 50 µM unlabeled or isotope-labeled aldoximes. For IAOx feeding, unlabeled or deuterium-labeled IAOx (D_5_-IAOx, with deuterium incorporated into the indole ring) was used. For PAOx feeding, unlabeled or deuterium-labeled PAOx (D_5_-PAOx, with deuterium incorporated into the benzyl ring) was used. For benzyl cyanide feeding, benzyl cyanide-α-^13^C was applied. After 24 h of incubation, seedlings were removed from the solution, rinsed with water, gently blotted dry, weighed, frozen in liquid nitrogen, and stored at −80 °C until extraction. A solution containing 0.005% (v/v) Triton X-100 without aldoxime was used as a mock control. Four biological replicates were included for each treatment. Labeled and unlabeled IAOx and PAOx were synthesized as previously described ^22^. Benzyl cyanide-α-^13^C was purchased from MilliporeSigma (Burlington, MA, USA).

### Detection and Quantification of Aldoximes, Auxins, and Benzyl Cyanide

Metabolites were extracted per mg fresh weight in 50 µL cold sodium phosphate buffer (50 mM, pH 7.0) containing sodium diethyldithiocarbamate and internal standards ([^13^C_6_]-IAA and [^13^C_6_]-PAA). Samples were incubated at 4 °C for 40 min with shaking, centrifuged (13,000 × g, 15 min, 4 °C), acidified to pH 2.7, and purified using Oasis™ HLB solid-phase extraction columns. Eluates (80% methanol) were dried and stored at −20 °C. For LC-MS analysis, samples were resuspended in water and analyzed using a Thermo Scientific Vanquish Horizon UHPLC coupled to a TSQ Altis Triple Quadrupole MS/MS with an Eclipse Plus C18 column (2.1 × 50 mm, 1.8 µm). For IAA, IAOx, and PAOx, positive ionization mode (4800 V) was used with 0.1% formic acid in water (A) and acetonitrile (B) under a 0–95% B gradient (4 min, 0.4 mL min⁻¹). For PAA, negative ionization mode (−4500 V) was used under identical chromatographic conditions. MRM transitions were: IAA (175.983→130.071), [^13^C_6_]-IAA (182.091→136), D_5_-IAA (181.102→134.083), IAOx (175.087→158), PAOx (136.076→119), PAA (135.045→91), and [^13^C_6_]-PAA (141.065→97).

For GC-MS analysis of PAA and benzyl cyanide, flash-frozen tissues were extracted in H_2_O:1-propanol: HCl (1:2:0.005), homogenized, and further extracted with methylene chloride. For unlabeled PAA quantification, a ^13^C_6_-PAA internal standard was added to the extract. Following centrifugation, the methylene chloride:1-propanol layer was derivatized with trimethylsilyldiazomethane, quenched with acetic acid in hexane, and volatile compounds were collected by vapor phase extraction. Volatiles were eluted in methylene chloride and analyzed using GC-MS with chemical ionization with isobutane. Derivatized PAA (methyl phenylacetate) labeled with C_13_ on the α-carbon (^1^C_13_-PAA) was monitored using extracted ion chromatography (EIC) at 152 m/z, unlabeled PAA at 151 m/z, D_5_-PAA at 156 m/z, and the internal standard ^13^C_6_-PAA at 157 m/z. Unlabeled BC was monitored at EIC 118 m/z and D_5_-BC at 123 m/z. Compound retention times and mass were confirmed using authentic standards.

### Generation and Validation of Inducible Tomato Lines

The *pOpON:AtCYP79A2* (*pOpA2*) and *pOpON:AtCYP79B2* (*pOpB2*) constructs were generated by Gateway cloning using entry vectors containing *AtCYP79A2* or *AtCYP79B2* and the *pOpON* vector ^39,40^, and subsequently introduced into *Agrobacterium tumefaciens* strain GV3101. Tomato transformation was performed using cotyledon explants from 10-day-old Micro-Tom seedlings grown on half-strength MS medium. Detached cotyledons were co-cultivated with *Agrobacterium* for 5 min and then transferred to callus induction medium for 2 weeks, followed by shoot induction medium for 2 weeks, and subsequently to root induction medium until root formation. Regenerated plantlets were transferred to soil. All culture media were based on MS salts supplemented with sucrose, plant growth regulators, and selection antibiotics. Callus induction medium contained 2 mg L^-1^ zeatin, 300 mg L^-1^ timentin, and 1.5 mg L^-1^ kanamycin. Shoot induction medium contained 1 mg L^-1^ zeatin, 300 mg L^-1^ timentin, and 1.5 mg L^-1^ kanamycin. Root induction medium contained 1 mg L^-1^ IAA, 200 mg L^-1^ timentin, and 1.5 mg L^-1^ kanamycin. Homozygous single-insertion T3 lines were identified based on GUS staining segregation ratios in the T2 and T3 generations. Inducible expression of *AtCYP79A2* and *AtCYP79B2* was confirmed by dexamethasone (dex) treatment. Dex was dissolved in dimethyl sulfoxide (DMSO) and diluted to 20 µM in water containing 0.02% (v/v) Tween-20. The mock solution contained 0.02% (v/v) DMSO and 0.02% (v/v) Tween-20 without dex. For GUS assays, cotyledons were detached and incubated in dex or mock solution overnight prior to staining. Tissues were incubated in GUS staining solution at 37 °C for at least 4 h, then destained in 70% ethanol at 37 °C. The GUS staining solution consisted of 100 mM sodium phosphate buffer (pH 7.2), 0.5 mM potassium ferricyanide, 0.5 mM potassium ferrocyanide, 0.3% (v/v) Triton X-100, and 1.9 mM X-Gluc.

### HPLC Analysis of Soluble Metabolites

Soluble metabolites were extracted from plant tissues in 50% (v/v) methanol at 65 °C for 2 h at a tissue concentration of 200 mg mL^-1^. Samples were centrifuged at 13,000 rpm for 10 min, and the supernatants were collected for HPLC analysis. Aliquots (10 µL) were analyzed using an UltiMate 3000 HPLC system (Thermo Fisher Scientific, Waltham, MA, USA) equipped with a diode array detector and a 10 °C autosampler. Metabolites were separated on an Acclaim™ RSLC120 C18 column (100 × 3 mm, 2.2 µm) using a mobile phase consisting of solvent A (0.1% (v/v) formic acid in water) and solvent B (acetonitrile) with a linear gradient from 3% to 95% solvent B over 20 min. The flow rate was 0.7 mL min^-1^, and the column temperature was maintained at 40 °C. Chlorogenic acid and rutin were quantified based on peak areas at 328 nm and 345 nm, respectively, using authentic standards (rutin, Sigma-Aldrich, St. Louis, MO, USA; chlorogenic acid, TCI, Portland, OR, USA). Three biological replicates were analyzed.

### Anthocyanin Measurement

Hypocotyls from dex-treated inducible lines or their mock-treated counterparts were collected and immersed in 1 mL of extraction buffer (18% 1-propanol, 1% HCl, and 81% water). Samples were boiled for 3 min and incubated in the dark at room temperature overnight. After centrifugation at 13,000 rpm for 10 min, the absorbance of the supernatant was measured at 535 nm and 650 nm using a spectrophotometer. Anthocyanin content was calculated as (A535 − A650) per g fresh weight.

### RNA Sequencing and Differentially Expressed Genes Analysis

For the *pOpA2* lines, seedlings grown on half-strength MS medium and treated with 20 μM dex or mock for two weeks were used for RNA extraction. For the *pOpB2* lines, the third and fourth leaves were collected from three-week-old soil-grown plants after seven days of dex spraying. Total RNA was extracted with TRIzol. Paired-end, unstranded, 150-bp sequencing of these total RNA samples was performed on an Illumina NovaSeq 6000 system. Raw reads were trimmed and filtered using AdapterRemoval v2. Trimmed reads were aligned to the tomato reference genome (ITAG 4.0) using HISAT2 v2.2.1. Read counts were generated with the featureCounts function from Subread v2.0.0, with fractional counting enabled for overlapping and multimapping reads. Differential expression analysis was performed using the DESeq2 package in the R statistical environment. To filter out lowly expressed genes, only genes with a total read count of 10 or more across all samples were retained. Log_2_ fold changes were shrunk using the normal prior. Significance was tested using the Wald test, and differentially expressed genes (DEGs) were defined using Benjamini–Hochberg-adjusted P-value cutoffs as indicated in the corresponding figures and figure legends. Gene Ontology (GO) biological process enrichment analyses were subsequently conducted on the identified DEGs using the clusterProfiler R package, with an adjusted P-value cutoff of 0.05.

### PAL Activity Assay

PAL activity was determined with minor modifications to a previously reported method ^27^. Frozen tomato tissues were homogenized using a Benchmark BeadBlaster 24 homogenizer (Benchmark Scientific, Sayreville, NJ, USA), and the resulting powder was extracted with buffer containing 0.1 M Tris-HCl (pH 8.3), 10% (v/v) glycerol, and 5 mM DTT. Total soluble protein concentration was measured using Bradford Reagent (Sigma-Aldrich, St. Louis, MO, USA) according to the manufacturer’s protocol. For the enzyme assay, 150 μL of crude protein extract was combined with 400 μL of reaction buffer containing 5 mM L-phenylalanine and incubated at 37 °C for 90 min. The reaction was stopped by adding 40 μL of 30% (v/v) acetic acid. Trans-cinnamic acid produced by PAL activity was extracted with 600 μL of ethyl acetate, dried using an Eppendorf Vacufuge Plus (Eppendorf, Hamburg, Germany), and resuspended in 100 μL of 50% (v/v) methanol. A 10 μL aliquot was subjected to HPLC analysis. Separation was performed using solvent A (0.1% (v/v) formic acid in water) and solvent B (acetonitrile) at a flow rate of 0.7 mL min^-1^, with the column maintained at 40 °C. Solvent B was increased from 12% to 30% over 2.6 min, from 30% to 95% over 4 min, and then held at 95% for 3 min. Trans-cinnamic acid was monitored at 270 nm and quantified based on the peak area using an authentic trans-cinnamic acid standard (Sigma-Aldrich). PAL activity was normalized to total soluble protein content.

## Acknowledgments

This work was supported by the National Science Foundation CAREER Grant (IOS-2142898) and the USDA National Institute of Food and Agriculture, Research Capacity Fund (Hatch) project 7004334 to JK, the United States Department of Agriculture-Agricultural Research Service project (6036-11210-002-000-D) to AKB, and NIH R35 grant (GM128742) to YD.

## Author Contributions

H.Z., D.S., and J.K. designed the research project. H.Z., D.S., E.T., K.H.C., A.S., D.L., Y.D., and A.K.B. performed the experiments and analyses. H.Z. and D.S. analyzed the data, generated the figures, and wrote the original draft. H.Z., D.S., and J.K. revised and edited the manuscript. Y.D., A.K.B., and J.K. acquired funding. J.K. supervised the project. All authors read and approved the final manuscript.

## Data availability

The raw RNA-seq reads generated in this study have been deposited in the NCBI Sequence Read Archive (SRA) under BioProject accession number PRJNA1460898.

## Supplementary Figures

**Supplementary Figure S1.** Phylogenetic analysis of tomato CYPs and the characterized CYP79s from other plant species.

**Supplementary Figure S2.** Amino acid sequence alignment of tomato CYP79 homologs and characterized CYP79 enzymes from other plant species.

**Supplementary Figure S3.** LC-MS/MS product ion spectra identifying aliphatic aldoximes produced by SlCYP79 enzymes.

**Supplementary Figure S4.** Pairwise amino acid sequence identity matrix of plant nitrilase homologs.

**Supplementary Figure S5.** Phenylalanine supplementation fails to restore phenylpropanoid accumulation in dex-treated *pOpA2* lines.

**Supplementary Figure S6.** Transcriptional activation of auxin-related genes in aldoxime-overproducing lines.

**Supplementary Figure S1.**
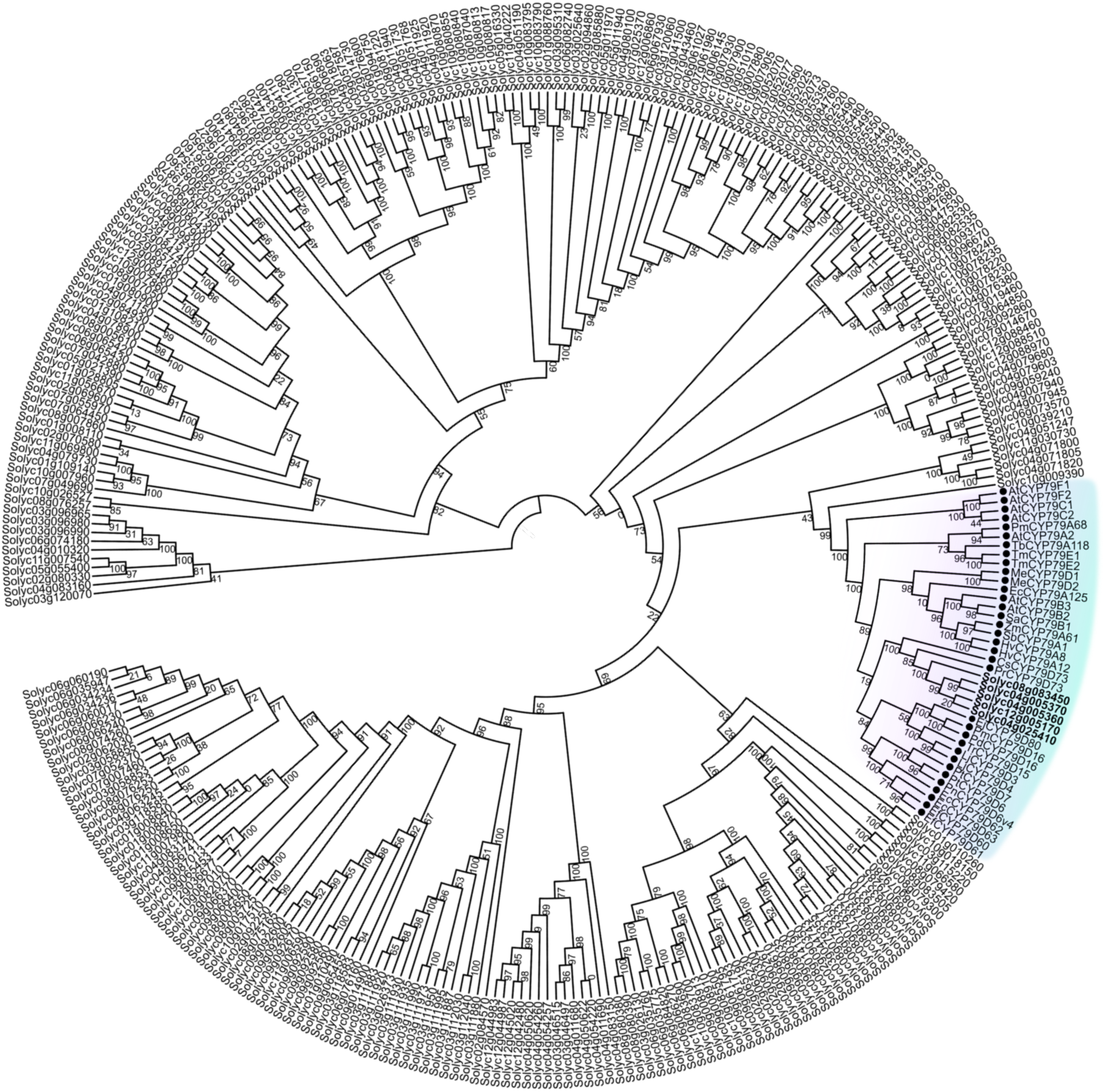
Phylogenetic analysis of tomato CYPs and the characterized CYP79s from other plant species. The tree includes five tomato CYP79 homologs and characterized CYP79 enzymes from other plant species. A light cyan shadow highlights the CYP79 genes. The rooted tree was inferred with the Maximum Likelihood method with 1000 bootstrap replicates. Species abbreviations: Sl, *Solanum lycopersicum*; At, *Arabidopsis thaliana*; Zm, *Zea mays*; Sb, *Sorghum bicolor*; Ec, *Erythroxylum coca*; Ef, *Erythroxylum fischeri*; Tb, *Taxus baccata*; Pr, *Plumeria rubra*; Ej, *Eriobotrya japonica*; Pm, *Prunus mume*; Pt, *Populus trichocarpa*; Pn, *Populus nigra*; Lj, *Lotus japonicus*; Me, *Manihot esculenta*; Tr, *Trifolium repens*; Hv, *Hordeum vulgare*; Pd, *Prunus dulcis*; Cs, *Camellia sinensis*; Sa, *Sinapis alba*; EcCYP79A125, *Eucalyptus cladocalyx*; Tm, *Triglochin maritima*.

**Supplementary Figure S2.**
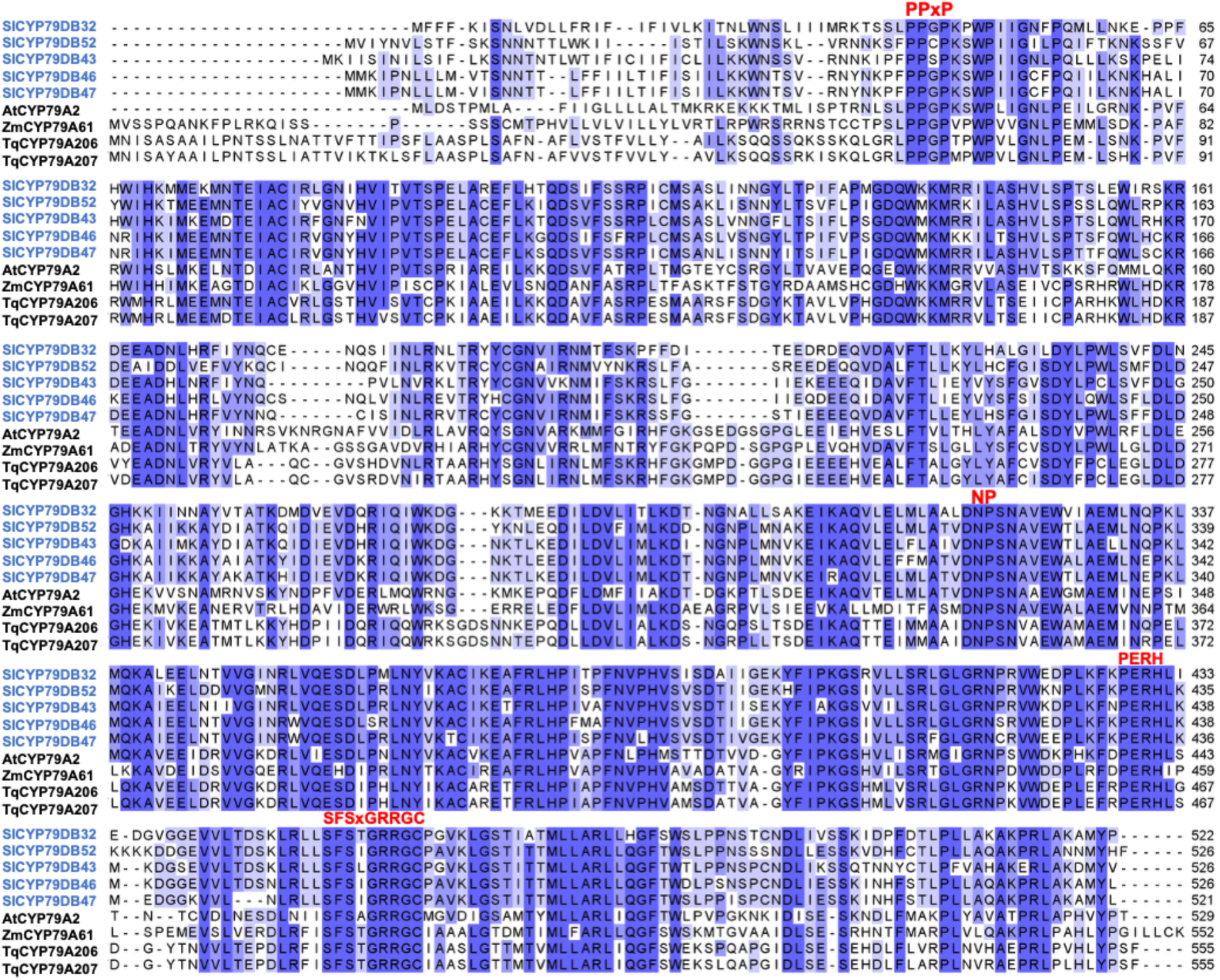
Amino acid sequence alignment of tomato CYP79 homologs and characterized CYP79 enzymes from other plant species. The alignment was generated using Clustal Omega (https://www.ebi.ac.uk/jdispatcher/msa/clustalo). Dark blue shading indicates highly conserved residues that are identical or similar across most sequences. Light blue shading indicates moderately conserved residues that are conserved in some sequences but vary in others. Red labels indicate sequence features discussed in the text, including the proline-rich PPGP/PPXP region, the NP motif, the PERH motif, and the heme-binding region containing the conserved cysteine residue, based on previous reports ^34–36^. Sl, *Solanum lycopersicum*; At, *Arabidopsis thaliana*; Zm, *Zea mays*; Tq, *Tococa quadrialata*.

**Supplementary Figure S3.**
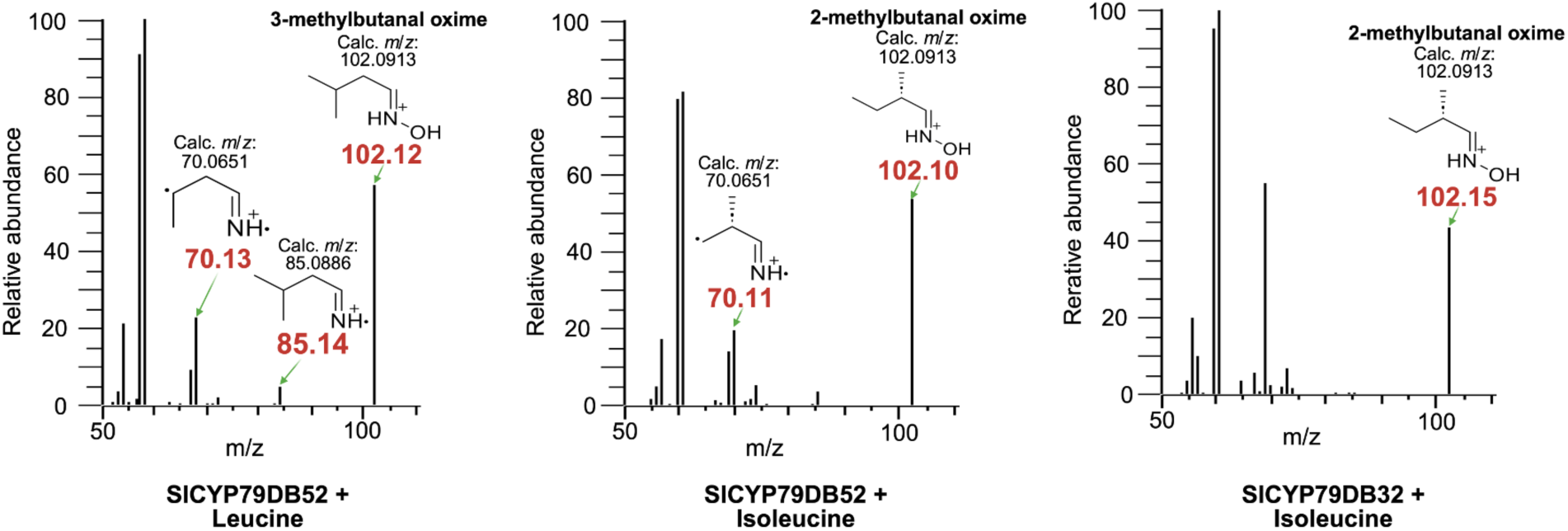
LC-MS/MS product ion spectra identifying aliphatic aldoximes produced by SlCYP79 enzymes. MS/MS fragmentation spectra corresponding to the reaction products shown in Figure 1E. The left panel displays the spectrum of the 3-methylbutanal oxime peak detected in the reaction of SlCYP79DB52 with leucine. The middle panel shows the spectrum of the 2-methylbutanal oxime peak detected in the reaction of SlCYP79DB52 with isoleucine. The right panel shows the spectrum of the 2-methylbutanal oxime peak detected in the reaction of SlCYP79DB32 with isoleucine. All panels display the relative abundance of ions plotted against their mass-to-charge ratio (m/z). Proposed chemical structures for the parent ion ([M+H]^+^, m/z ≈ 102) and key fragment ions are illustrated within each panel. Red numbers indicate experimentally observed m/z values, whereas black numbers represent calculated theoretical m/z values.

**Supplementary Figure S4.**
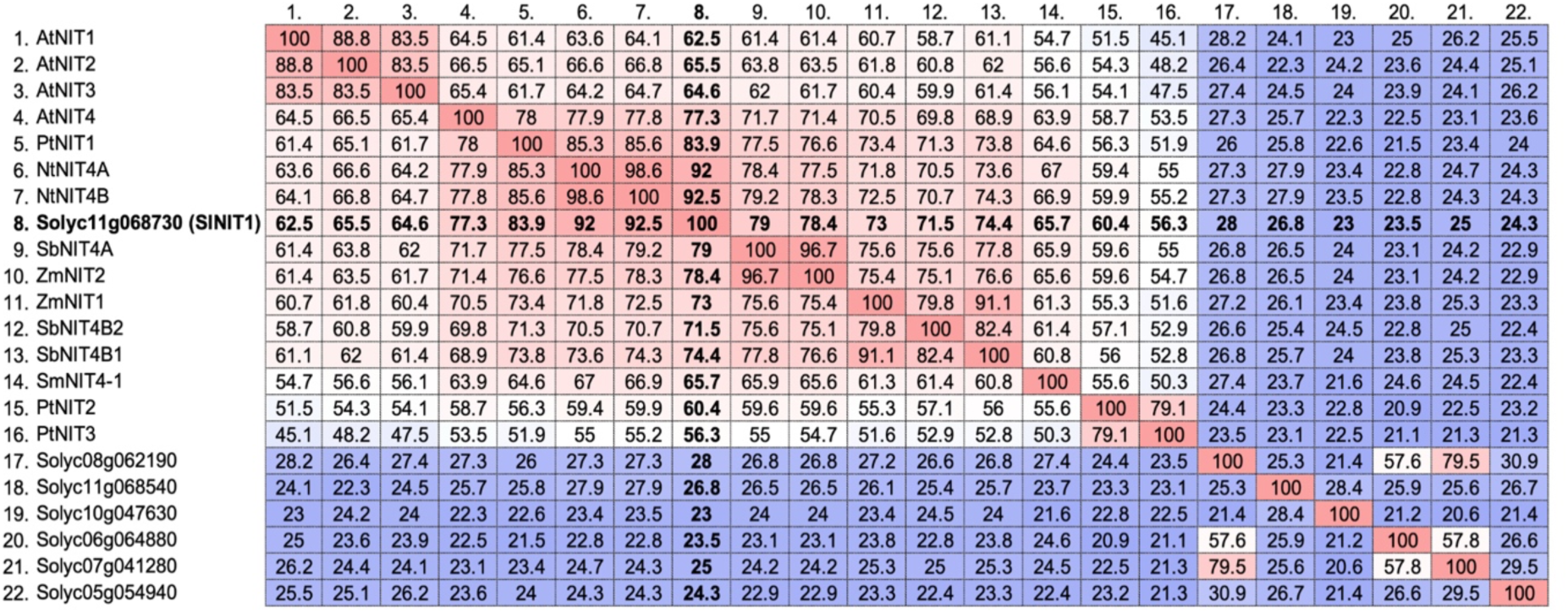
Pairwise amino acid sequence identity matrix of plant nitrilase homologs. The matrix displays the percentage identity between full-length amino acid sequences of NIT proteins from tomato (*Solanum lycopersicum*) and other characterized plant species used in the phylogenetic analysis shown in Fig. 3D. Sequence alignments and pairwise identity calculations were performed using the Clustal Omega multiple sequence alignment tool (EMBL-EBI). The numbers within the cells represent the percent identity, and the color scale indicates the degree of similarity, ranging from blue (low identity) to red (high identity). Species abbreviations: *Sl*, *Solanum lycopersicum*; *At*, *Arabidopsis thaliana*; *Zm*, *Zea mays*; *Sb*, *Sorghum bicolor*; *Nt*, *Nicotiana tabacum*; *Pt*, *Populus trichocarpa*; *Sm*, *Selaginella moellendorffii*.

**Supplementary Figure S5.**
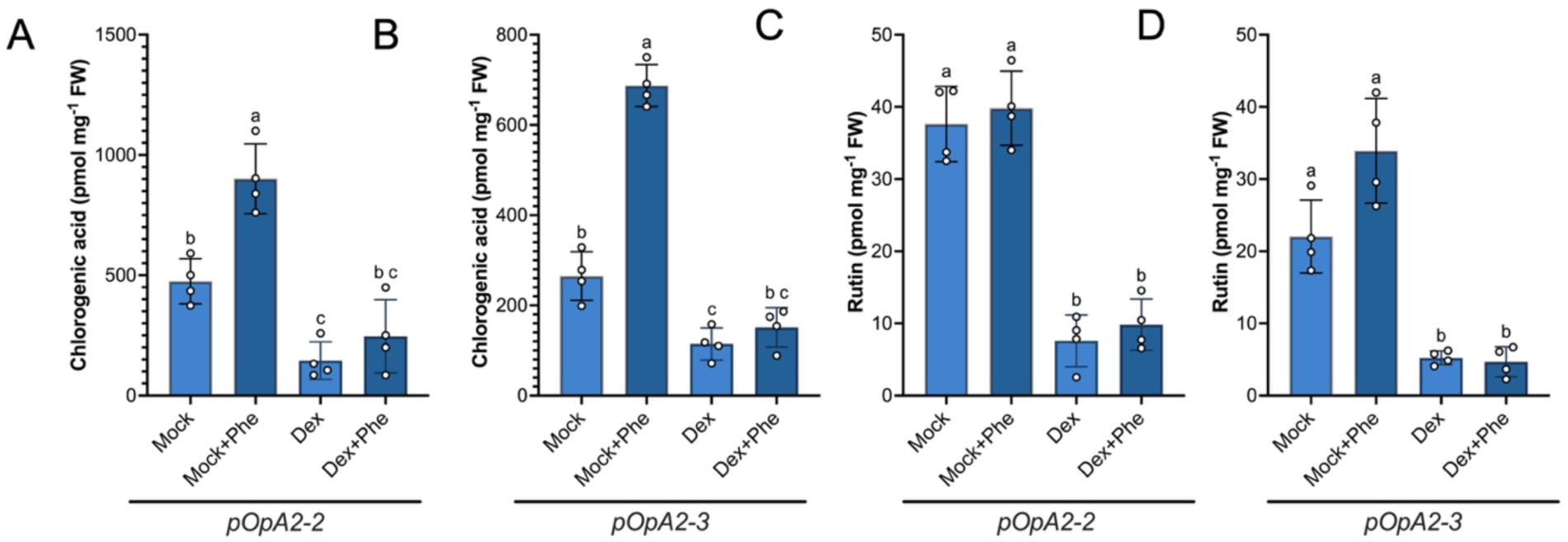
Phenylalanine supplementation fails to restore phenylpropanoid accumulation in dex-treated *pOpA2* lines. (A, B) Chlorogenic acid levels in *pOpA2-2* (A) and *pOpA2-3* (B) seedlings. (C, D) Rutin levels in *pOpA2-2* (C) and *pOpA2-3* (D) seedlings. Seedlings grown on half-strength MS medium were treated for two weeks with mock, mock + 100 μM phenylalanine (Phe), 20 μM dexamethasone (dex), or 20 μM dex + 100 μM Phe. Exogenous Phe supplementation did not restore the reduced chlorogenic acid or rutin levels in dex-treated plants. Data represent the mean ± SD of four biological replicates, with each replicate containing at least two seedlings. Different lowercase letters indicate significant differences among treatments within each panel, as determined by one-way ANOVA followed by Tukey’s multiple comparisons test (P < 0.05).

**Supplementary Figure S6.**
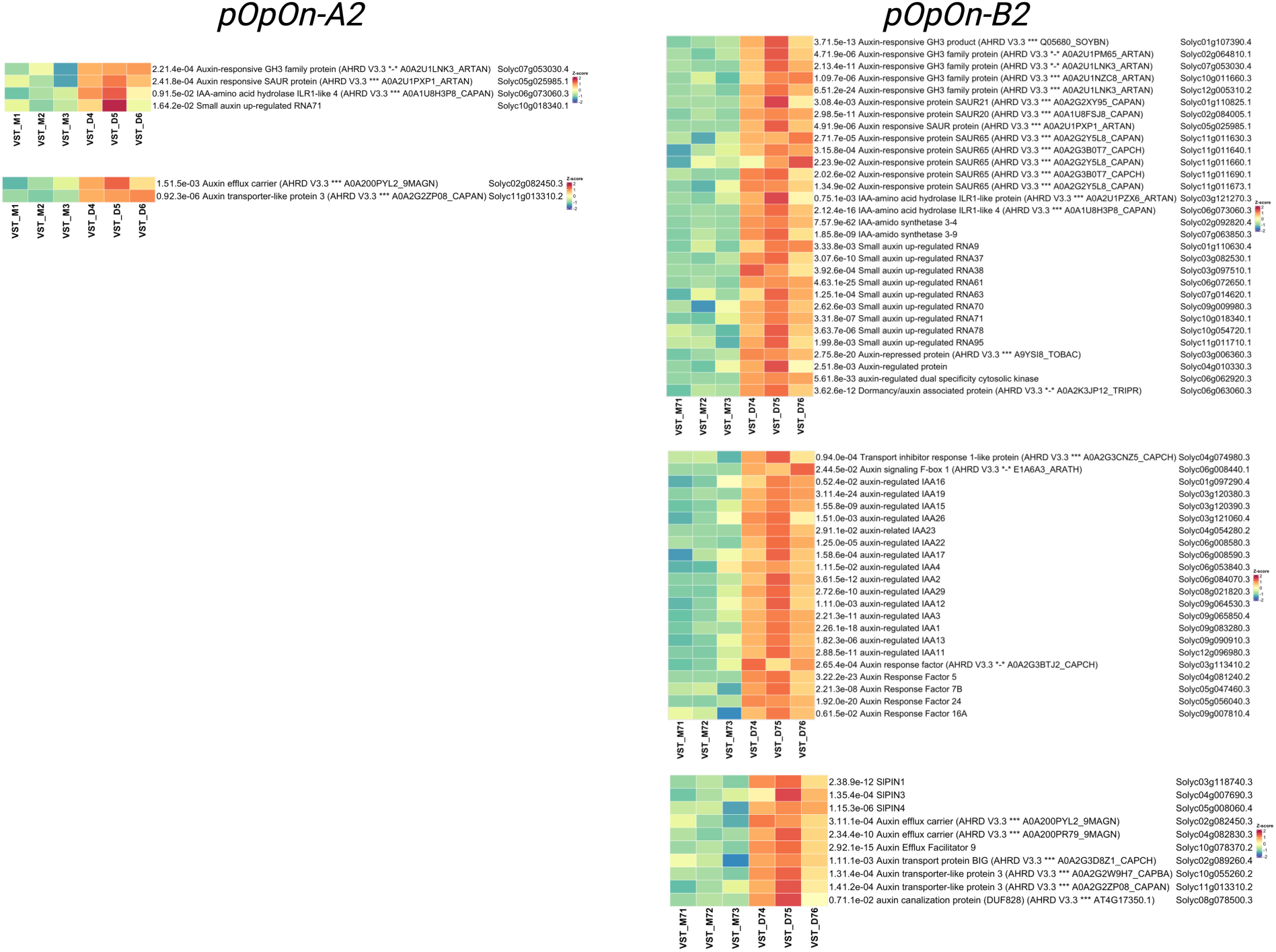
Transcriptional activation of auxin-related genes in aldoxime-overproducing lines. Heatmap analysis of differentially expressed genes associated with auxin biology in the *pOpA2* and *pOpB2* lines. These auxin-related genes were extracted from the significant differentially expressed gene datasets identified using the DESeq2 pipeline. Differential expression was determined using an adjusted P-value cutoff of 0.05. The color scale represents row-scaled Z-scores of transcripts per million values, where red indicates higher relative expression and blue or green indicates lower relative expression. Gene descriptions, adjusted P-values, and Solyc IDs are provided to the right of each heatmap row.

